# A robust modular automated neuroimaging pipeline for model inputs to TheVirtualBrain

**DOI:** 10.1101/2022.02.24.481836

**Authors:** Noah Frazier-Logue, Justin Wang, Zheng Wang, Devin Sodums, Anisha Khosla, Alexandria Samson, Anthony R. McIntosh, Kelly Shen

## Abstract

TheVirtualBrain, an open-source platform for large-scale network modelling, can be personalized to an individual using a wide range of neuroimaging modalities. With the growing number and scale of neuroimaging data sharing initiatives of both healthy and clinical populations comes an opportunity to create large and heterogeneous sets of dynamic network models to better understand individual differences in network dynamics and their impact on brain health. Here we present TheVirtualBrain-UK Biobank pipeline, a robust, automated and open-source brain image processing solution to address the expanding scope of TheVirtualBrain project. Our pipeline generates connectome-based modelling inputs compatible for use with TheVirtualBrain. We leverage the existing multimodal MRI processing pipeline from the UK Biobank made for use with a variety of brain imaging modalities. We add various features and changes to the original UK Biobank implementation specifically for informing large-scale network models, including user-defined parcellations for the construction of matching whole-brain functional and structural connectomes. Changes also include detailed reports for quality control of all modalities, a streamlined installation process, modular software packaging, updated software versions, and support for various publicly available datasets. The pipeline has been tested on both healthy and clinical populations and is robust to the morphological changes observed in aging and dementia. In this paper, we describe these and other pipeline additions and modifications in detail, as well as how this pipeline fits into the TheVirtualBrain ecosystem.

## 1 Introduction

Neuroimaging data sharing initiatives have expanded substantially in the last decade. Multimodal data collection initiatives like the Human Connectome Project (HCP; Van Essen et al., 2013)), UK Biobank (Sudlow et al., 2015), and Alzheimer’s Disease Neuroimaging Initiative (ADNI; Mueller et al., 2005), among others, allow for promising new avenues of neuroscientific research that connect different scales of measurement across large samples. While many efforts are being made to analyze these large datasets to better understand the inner workings of the brain and, specific to neurological disorders, identify effective biomarkers of disease, their potential for creating large and heterogeneous sets of personalized generative models is not yet fully realized. TheVirtualBrain (TVB) is an open source software platform for large-scale network modelling (Sanz Leon et al., 2013; Sanz-Leon et al., 2015), where models can be personalized to an individual using a wide range of neuroimaging modalities. Creating personalized models in TVB from large multimodal neuroimaging datasets will allow us to not only better understand individual differences in network dynamics but also allow for the interrogation of mechanisms of disease across large and heterogeneous samples.

For modelling large-scale brain networks, TVB requires, as input, a structural connectivity matrix that represents the anatomical wiring of the brain. In humans, this is often derived from anatomical (T1w) and diffusion-weighted magnetic resonance imaging (dMRI) tractography and specified as the long-range connections between brain regions of interest (ROIs). Optional inputs for TVB models include the cortical surface for surface-based models (e.g., Spiegler et al., 2016), and functional data (e.g., BOLD-fMRI responses, M/EEG activity, functional connectivity) for model input (e.g., Schirner et al., 2018) or parameter fitting (e.g., Shen et al., 2019a), parcellated into the same ROIs as the structural connectivity. A software pipeline for processing large datasets for TVB, then, would ideally be automated and able to preprocess multiple imaging modalities into a set of matching parcellated inputs for TVB. Existing popular MRI processing pipelines include fmriPrep for anatomical and fMRI data (Esteban et al., 2019), and HCP’s Minimal Preprocessing Pipeline for anatomical, fMRI and dMRI data (Glasser et al., 2013). HCP’s pipeline is especially well suited for higher resolution images and relies on the FreeSurfer software package (Fischl, 2012) for working with the cortical surface. An existing empirical data processing pipeline already exists for processing anatomical, fMRI and dMRI data for TVB inputs, and also relies on FreeSurfer-generated surfaces (Schirner et al., 2015).

Data re-use of publicly-available datasets offers great promise for improving both accessibility and replicability. Within the scope of connectome-based modelling, these data also present the opportunity to generate models that capture a population-level understanding that no single empirical dataset can offer. However, considerations for data processing and analysis of data acquired using older protocols and in special populations are warranted. For example, a user may wish to avoid the projection of lower resolution data (e.g., fMRI) to cortical surface vertices (Alfaro-Almagro et al., 2018). With data from aging and clinical populations, FreeSurfer tissue-class segmentations can also be inaccurate and may require manual intervention (McCarthy et al., 2015; Henschel et al., 2020; Srinivasan et al., 2020), something that is not reasonably feasible with large samples. Moreover, with automated processing, a quality control (QC) workflow that detects processing inaccuracies is also needed. This is especially important for aging and clinical datasets where inaccuracies in preprocessing MRI data are common due to differences in brain morphology and image contrast. The HCP pipeline can be used with an fMRI QC pipeline that computes summary statistics to capture signal quality and subject motion of fMRI scans (Marcus et al., 2013). QC of other imaging modalities (e.g., T1w, dMRI) processed with the HCP pipeline relies on extensive manual inspection of raw and processed images. MRIQC (Esteban et al., 2017) is an fMRIPrep-compatible software package that computes image-based metrics for raw or minimally-processed T1w and fMRI data. It outputs a set of html-based reports of the individual and group-wise summary metrics to allow identification of outlier images. MRIQC also offers an automated pass-fail classification of T1w images. These existing tools, however, do not allow for identification for common preprocessing errors such as poor tissue-class segmentation, and poor registrations to templates and across modalities. Often, these errors are detected via detailed manual QC but the visual inspection of hundreds to thousands of subject’s processed multi-modal data derivatives is unfeasible and a streamlined QC workflow at the scale of such large datasets is needed.

The UK Biobank offers an alternative multi-modal MRI (anatomical, fMRI, dMRI, susceptibility-weighted MRI) processing pipeline that mostly relies on tools from the FMRIB Software Library (FSL; Jenkinson et al., 2012) and maintains images in volumetric space. The pipeline is fully automated, built to process the very large and longitudinal UK Biobank sample of aging individuals. It generates a number of image-based metrics of raw and processed intermediates, mostly from their structural preprocessing sub-pipeline. Referred to as “Imaging-Derived Phenotypes”, these metrics were used for automated QC of the large UK Biobank aging sample. Here, we describe an extension of the UK Biobank pipeline that addresses the expanding scope of TheVirtualBrain project. The extension includes the generation of matched structural and functional connectivity data based on a user-defined brain parcellation, expanded capability for additional MRI modalities and manufacturers, additional preprocessing considerations for aging data (e.g., age-specific templates), an expanded number of image-based metrics for fMRI and dMRI, and the addition of new metrics for structural and functional connectivity. We have also developed an extensive new HTML-based QC report for quick assessment of raw, intermediate and processed outputs, and containerized the pipeline to maximize portability and ease of installation. The pipeline supports data from aging and neurodegenerative populations, and has been tested on a number of different datasets including multi-modal MRI data from the Cambridge Centre for Ageing and Neuroscience study (Cam-CAN; Taylor et al., 2017) as well as the ADNI3 study (Weiner et al., 2016). Finally, in keeping with TheVirtualBrain’s commitment to the FAIR guiding principles (Wilkinson et al., 2016) and open science practices, our pipeline is open source and compliant with the Brain Imaging Data Structure (BIDS) standard (Gorgolewski et al., 2016). Below, we describe the software and methodological modifications and additions we made to the original UK Biobank pipeline, highlight the new QC pipeline and HTML report, show some usage examples, and discuss future work and integrations with TheVirtualBrain.

## 2 Method

We refer to our pipeline as TheVirtualBrain-UK Biobank (or TVB-UKBB) pipeline. It is built from a fork of the UK Biobank pipeline (https://git.fmrib.ox.ac.uk/falmagro/UK_biobank_pipeline_v_1), which has been previously described (Alfaro-Almagro et al., 2018). The UK Biobank pipeline processes a variety of MRI modalities but, for the purposes of creating TVB inputs, we focused on modifying and extending the existing structural (T1w, T2 FLAIR), functional (resting-state, task), and diffusion-weighted MRI sub-pipelines. The processing of other MRI modalities (e.g., susceptibility-weighted imaging) in the TVB-UKBB pipeline remain unaltered and untested.

Figure 1 shows the general workflow of the whole pipeline, its sub-pipelines, and their outputs. The major output of the structural MRI pipeline is the user-defined parcellation registered to the subject’s T1w image. The registered parcellation is used by both the functional and diffusion MRI sub-pipelines to define ROIs for computing average regional timeseries and connectivity measures for TVB inputs. Following the completion of the functional and diffusion MRI sub-pipelines, an ‘IDP’ pipeline computes image-based metrics for all modalities. Finally, our newly developed QC pipeline generates a comprehensive HTML-based report for manual quality assurance procedures.

**Figure 1.**
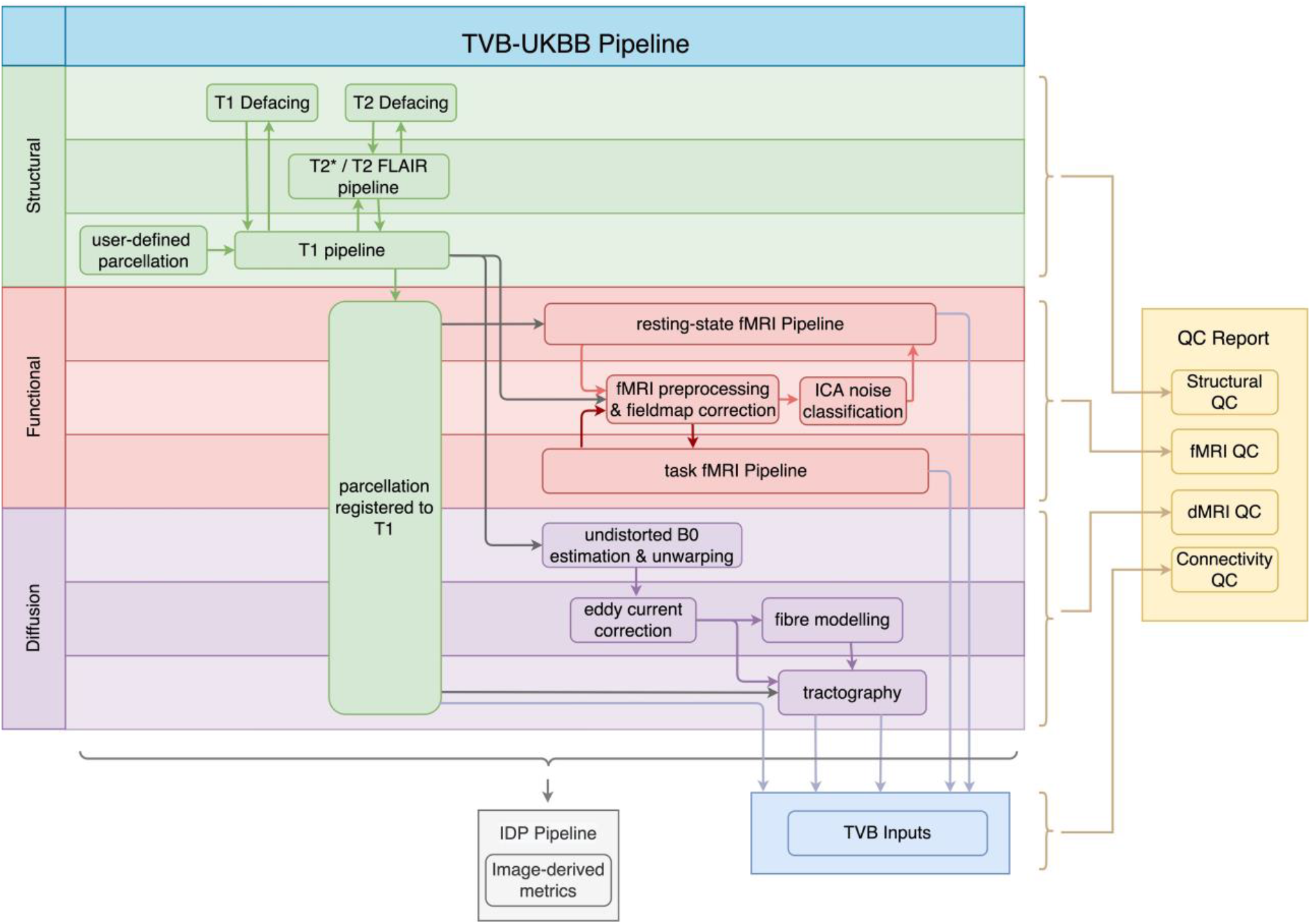
General workflow of the TVB-UKBB pipeline. The main imaging sub-pipelines of interest for the current paper are shown (structural in green, functional in red, and diffusion in purple). A TVB-compatible .zip file (TVB Inputs) is created from the relevant outputs of the imaging sub-pipelines. The ‘IDP Pipeline’ collects image-based metrics from raw, intermediate and process outputs across imaging sub-pipelines and make them available for analysis. The final step of the pipeline is the generation of the QC report.

### 2.1 Structural sub-pipeline

Our pipeline largely retains the structural (T1w, T2 FLAIR) preprocessing steps from the UK Biobank pipeline (Alfaro-Almagro et al., 2018). These include brain extraction and nonlinear registration to the MNI152 standard-space T1 template, defacing, bias correction and tissue-class segmentation (Figure 2). Processing of T2* images (brain extraction, registration to MNI152 and T1w, bias correction) has been added. Other major modifications and additions to the structural sub-pipeline are outlined below.

**Figure 2:**
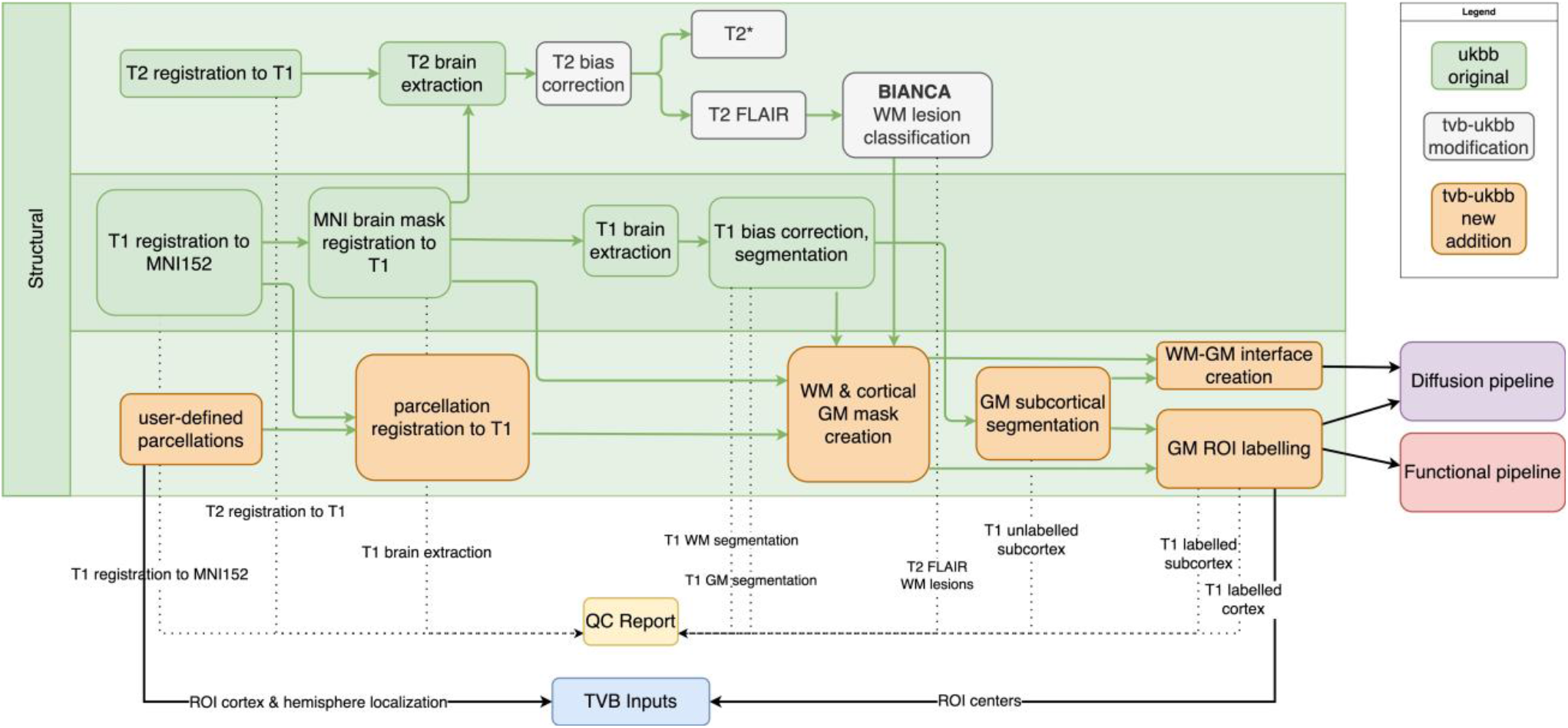
Structural sub-pipeline workflow. Original components of the UK Biobank pipeline with few or no modifications are in green; pipeline components with major changes or additions are indicated in white; and new components are indicated in orange. Dotted lines indicate components that are included in the QC report. Black lines indicate components that are used downstream by other sub-pipelines or included in ‘TVB Inputs’. GM: grey matter, WM: white matter

#### 2.1.1 Parcellation

To support connectome-based modelling in TVB, our additions to the structural sub-pipeline allow users to create connectomes from T1w, dMRI and resting-state fMRI data by specifying a brain parcellation of their choice. Currently, our pipeline supports parcellations defined on the MNI152 1mm template. For ease, we include three different parcellations in our repository. Two are combinations of the Schaefer cortical (Schaefer et al., 2018) with either the Tian subcortical (Tian et al., 2020) or Harvard-Oxford subcortical (Frazier et al., 2005) parcellation and the third is the Regional Map parcellation (Bezgin et al., 2017). The Schaefer-Tian parcellation is offered at three different scales of granularity and, if the user wishes, other scales can be created from the parcellations shared on the respective GitHub repositories. A tab-separated look-up table for the parcellation that specifies image labels and label names is required. The parcellation is registered to the T1w image using the warps from the nonlinear registration of the template to T1w.

#### 2.1.2 Segmentation

In both healthy older adult and neurodegenerative samples, accurate tissue classification using T1w images is hindered by decreasing image contrast with age (Bansal et al., 2013). Additional difficulties in T1w tissue classification arise from the presence of white matter pathology, where white matter lesions become misclassified as grey matter (Levy-Cooperman et al., 2008). Since tissue classification is a vital part to defining accurate ROIs for both structural and functional connectivity, we have implemented a number of modifications to the segmentation procedure to improve ROI assignments. We derive an initial image segmentation following the UK Biobank’s procedure using FSL’s FAST toolbox. We then refine the grey matter subcortical segmentation by adding the outputs of FSL’s FIRST toolbox (an object model-based segmentation and registration tool) to the grey matter mask.

To address inaccuracies in the grey matter mask due to the presence of WM pathology, we have implemented two alternative methods that may be used depending on available image modalities. The first method, if T2 FLAIR images are available, uses the outputs of the WM lesion classification (FSL’s BIANCA) to exclude any misclassified voxels from the grey matter mask and add them back to the white matter mask. The second method is an option for when T2 FLAIR images are not available. In these cases, we use age-specific image classes (Fillmore et al., 2015) as tissue priors. T1w images from adults aged 40 or over are registered to the template for their age decile (e.g., 40s, 50s, etc.) while subjects aged under 40 are registered to the FSL-distributed tissue priors. These template space-registered T1w images are then segmented using the set of matching age-specific priors. Segmented images are registered back to T1w space. Age-specific templates are provided up to the 80s decile. Subjects greater than 89 years are registered to the 80s decile template.

#### 2.1.3 Defining Regions of Interest for fMRI and dMRI sub-pipelines

The user-provided parcellation is registered to the T1w image and the grey matter mask is labelled with ROI indices. The labelled grey matter volume serves as input to the functional MRI sub-pipeline. The white and grey matter segmentations are both used to create the grey matter-white matter interface for dMRI tractography. This interface consists of voxels of white matter adjacent grey matter and, when labelled, will serve as the seed and target masks for tractography in the diffusion MRI sub-pipeline.

### 2.2 Functional MRI sub-pipeline

The fMRI sub-pipeline processes both resting-state- and task-fMRI data (Figure 3). The processing of both data types by the UK Biobank pipeline relies on FSL’s FEAT toolbox. As best practices for preprocessing of fMRI data are both dataset-dependent and constantly evolving (Uddin, 2017), the pipeline allows users flexibility on selecting the right preprocessing methods for their needs. Users may specify their preferences, which can include brain extraction, motion correction via realignment of fMRI images (MCFLIRT), slice timing correction, spatial smoothing, intensity normalization, and temporal filtering. Registration to the T1w image and MNI152 template is performed. For resting-state fMRI data, automated classification and removal of noise artefacts is performed using FMRIB’s ICA-based Xnoiseifier (FIX) (Griffanti et al., 2014).

**Figure 3:**
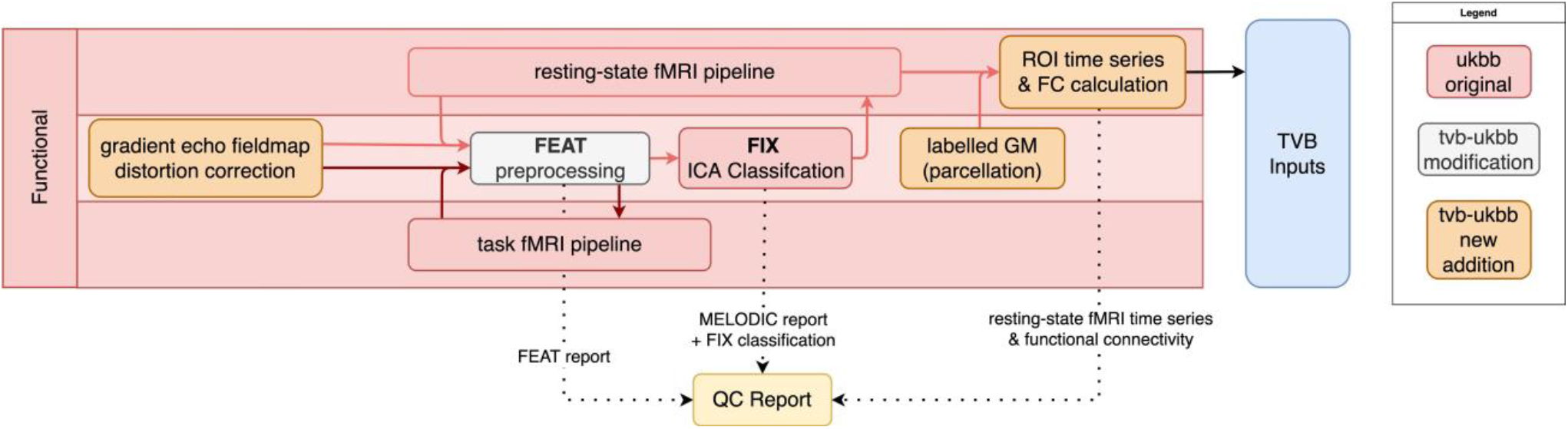
fMRI sub-pipeline workflow. Original components of the UKBB pipeline with few or no modifications shown in red; pipeline components with major changes or additions shown in white; and new components shown in orange. Dotted lines indicate components that are included in the QC report. Black lines indicate components that are included in the TVB Inputs.

We have modified the UK Biobank pipeline to now accept an arbitrary number of fMRI sessions. Other major additions and modifications are described below.

#### 2.2.1 Field map correction

The UK Biobank pipeline performs geometric distortion correction for the unwarping of EPI (e.g., fMRI and dMRI) images. This correction requires a reverse phase-encoded B0 dMRI image for estimating the field map, which is not always available. To support more “traditional” field map acquisitions for EPI distortion correction, such as those in the Cam-CAN dataset, we have implemented the option for dual echo-time gradient distortion correction using FSL’s FUGUE toolbox.

#### 2.2.2 Resting-state fMRI

We have updated the pipeline’s FIX version from 1.063 to 1.06.15. Although FMRIB provides a default trained-weights file, and we provide trained-weights files for both the ADNI3 and Cam-CAN datasets, the classifier performs best when trained with the user’s specific dataset. The most notable addition to resting-state fMRI processing is the replacement of group-ICA-based detection of resting-state networks with the parcellation of the resting-state fMRI data to accommodate connectome-based modelling. Following denoising, the parcellation output from the structural sub-pipeline (Figure 2) is registered to a reference resting-state fMRI volume and the average BOLD response across voxels is computed for all ROIs (i.e., ROI time series). The Pearson correlation coefficient between all ROI time series is also computed to obtain a measure of functional connectivity.

#### 2.2.3 Task-based fMRI

In our implementation of the fMRI sub-pipeline, task-based fMRI data are minimally preprocessed but not further analyzed. Users may choose to re-implement a GLM-based analysis using FEAT or, alternatively, they may take the preprocessed task-fMRI data and apply other analytic methods (e.g., Partial Least Squares; McIntosh and Lobaugh, 2004).

### 2.3 Diffusion sub-pipeline

Processing steps for diffusion imaging data that we have retained from the UK Biobank pipeline include correction of eddy currents and head motion (EDDY), diffusion tensor image fitting (DTIFIT) for tract-based analysis (TBSS), and multi-fibre orientation modelling (BEDPOSTX) (Figure 4). New features and additions to the diffusion sub-pipeline are described below.

**Figure 4:**
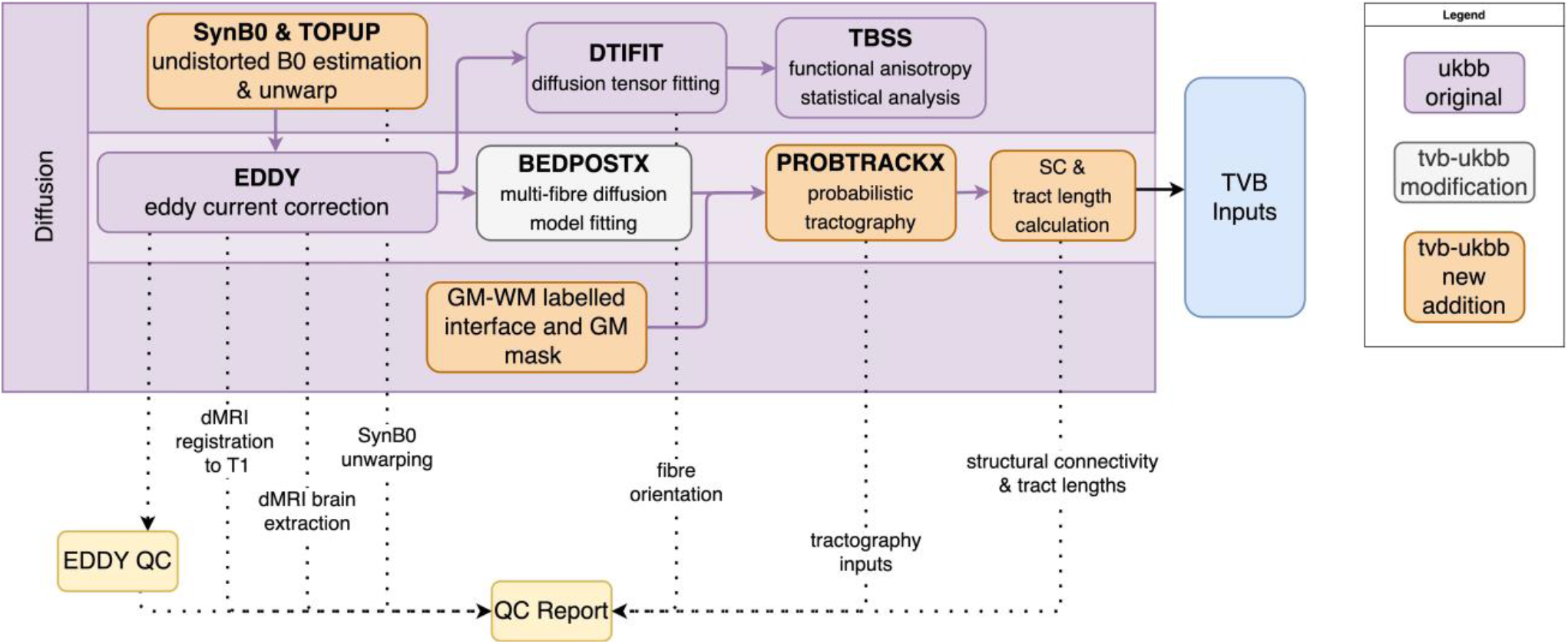
Diffusion sub-pipeline workflow. Original components of the UKBB pipeline with few or no modifications shown in purple; pipeline components with major changes or additions shown in white; and new components shown in orange. Dotted lines indicate components that are included in the QC report. Black lines indicate components that are included in the TVB Inputs.

#### 2.3.1 Distortion correction with synthesized B0

Our first addition to the diffusion sub-pipeline was the integration of B0 field estimation for unwarping data that lack reverse phase-encoded images using the Synb0-DisCo tool (Schilling et al., 2019). This tool uses a deep learning approach to create a synthetic undistorted B0 image from a T1w image. The synthetic undistorted B0 is used as input to FSL’s TOPUP toolbox for dMRI distortion correction. In our pipeline, users have the option to implement this tool to improve registrations between the T1w and dMRI images.

#### 2.3.2 Tractography

The other major addition to the dMRI sub-pipeline was the replacement of the UK Biobank tractography approach with one that takes as input the user-defined parcellation for connectome construction. In our approach, the grey matter-white matter labelled interface is registered to the distortion-corrected B0 image. This interface is used to define seed and target ROI masks. The grey matter mask is also registered to the B0 image and used as an exclusion mask. Probabilistic tractography is performed using FSL’s PROBTRACKX toolbox to generate a matrix of the streamlines between all ROIs. The structural connectivity ‘weights’ matrix is then computed by taking the streamlines matrix and dividing it by the total number of streamlines that were successfully sent from the seed ROIs. This weights matrix therefore encapsulates the probability of connection between all ROIs. ‘Distance’ matrices (i.e., estimated tract lengths) are also retained.

### 2.4 Compatibility with TheVirtualBrain

Our pipeline generates inputs for connectome-based modelling, with file formats that are directly compatible with TheVirtualBrain (TVB; thevirtualbrain.org) (Supplementary Figure 1). These include the structural connectivity weights and tract lengths matrices, as well as the ROI time series and functional connectivity matrix from resting-state fMRI scans. ROI location information such as hemisphere or subcortical localization and centroid coordinates are also included. Towards the end of the pipeline, these TVB-input files are given the appropriate file names, placed in the correct folder structure, and compressed into a zip file that can be accepted by TVB without further processing. This zip file can be found in the top-level directory for each processed subject.

### 2.5 Imaging Derived Phenotypes (IDPs)

The original UK Biobank pipeline generates various image-based metrics, or IDPs, for evaluating the characteristics of input images, pipeline processing outputs, and derivative files. These IDPs were intended to be a quantitative measure of the quality of processed subjects but mostly describe structural sub-pipeline processing and outputs. To better capture modalities of interest for connectome-based modelling, we have developed an additional 75 unique IDPs that describe fMRI and dMRI processing as well as connectivity outputs [Supplementary Table 1]. Notable examples include IDPs for assessing the alignment of various modalities to T1 space, the temporal signal-to-noise ratio (tSNR) in resting-state fMRI, and summary statistics for functional and structural connectivity. In conjunction with the original IDPs, these new metrics were developed for the purpose of flagging subjects whose outputs’ quality is poor, either due to acquisition errors, subject anomalies, or pipeline errors and insufficiencies.

We performed a manual QC of 140 (70 female, 70 male) Cam-CAN subjects using our QC reports (described below) to enable assessment of the utility of our newly developed IDPs for quantifying processing errors. The subjects were pseudorandomly selected, balanced for sex and 20 were chosen from each age decile to cover the entire age range of the dataset. Two experienced subject raters (DS, AK) scored the processing intermediates and outputs. These graders gave each subject a score along a 5-point scale for each modality (ranging from excellent [1] to poor [5]) and also gave each subject a pass/fail classification based on the integrity of the TVB inputs as a whole. A fuller description of the QC procedure and example QC report usage for the Cam-CAN data is presented in the Results section. We used a multivariate statistical approach, partial least squares analysis (Krishnan et al., 2011), to identify a set of latent variables that represent the maximal covariance between the QC ratings and the image-based metrics outputted from the pipeline. First, the covariance between the two sets of variables was computed. Singular value decomposition on this cross-block covariance was then performed to produce latent variables, each containing three elements: (1) a set of weighted “saliences” that describe a pattern of IDPs; (2) a design contrast of QC ratings that express their relation to the saliences, and (3) a scalar singular value that expresses the strength of the covariance. The mutually orthogonal latent variables are extracted in order of magnitude, with the first latent variable explaining the most covariance between the IDPs and QC ratings, the second LV the second most, and so on. We report the relative percentage of total cross-block covariance explained by each latent variable, where the sum of this percentage across all latent variables is 100. The statistical significance of each latent variable was assessed with permutation testing: 1000 permutations shuffled subjects’ QC ratings without replacement while maintaining their IDP assignments. This resulted in 1000 new covariance matrices which were each subjected to singular value decomposition to produce a null distribution of singular values. The reliability with which each

IDP expressed the differences across QC ratings was determined with bootstrapping: 500 bootstrap samples were created by resampling subjects with replacement within each rating class. This resulted in 500 new covariance matrices which were, again, subjected to singular value decomposition. The 500 saliences from the bootstrapped dataset were used to build a sampling distribution of the saliences from the original dataset. The bootstrap ratio for a given IDP was calculated by taking the ratio of the salience to its boostrap-estimated standard error. With the assumption that the bootstrap distribution is normal, the bootstrap ratio is akin to a Z-score and corresponding saliences were considered to be reliable if the absolute value of their bootstrap ratio was ≥ 2.

### 2.6 Quality Control Report

Typical manual QC requires users to manually search for NIFTI files, load them into visualizer GUIs like FSLeyes, and adjust various parameters for each overlay. To streamline these procedures, our pipeline generates a Quality Control (QC) Report for each subject. The QC sub-pipeline runs at the end of the TVB-UKBB pipeline and leverages derivative data to generate brain image overlays, data visualization plots, and summary tables. These assets are wrapped in an offline HTML page that can be compressed into a portable, small, and standalone archive using a script included in the pipeline. This standalone report may be viewed on any browser and requires no access to the original subject’s files.

Our QC Report allows users to view and interact with 17 preset key QC overlays immediately upon opening the HTML report. Our QC Report offers the ability to zoom, pan, switch between planes of view, inspect different analyses, and toggle visibility of layers in brain overlay images. These controls are also assigned to various hotkeys, allowing for browsing without a mouse and further expediting the QC process for more experienced users. Additionally, each brain overlay shows an array of 18 slices for each orientation, saving time typically spent seeking slices in visualization software. Especially when considering that multiple different overlays need to be generated for QC and certain overlays may need to be revisited more than once, our HTML Report can economize users’ time and effort in the QC process.

The QC Report features a page for each sub-pipeline and multiple analyses on each page, corresponding to various key steps of the sub pipeline. For instance, brain image overlays, generated using FSL’s FSLeyes and SLICER, are intended to offer users qualitative assessment of brain extraction, segmentation, registration, and labelling for multiple modalities (Figure 5). Data visualization plots are also included to simplify the verification of TVB-inputs. IDP tables offer a simple interface for accessing metrics and assessing the quality of a subject’s processing. Within these tables, rows of IDPs are colour-coded green or red (pass or fail) depending on their values relative to user-defined thresholds. A more detailed summary and explanation of QC analyses included in the report can be found in the Supplementary Material (Supplementary Tables 2-4). At the bottom of several QC Report pages, there are multiple file path links to the depicted overlay image as well as its source NIFTI image files. If more detailed investigation into a processed subject is required, then users have the option to load these files and perform QC with a NIFTI visualizer.

**Figure 5:**
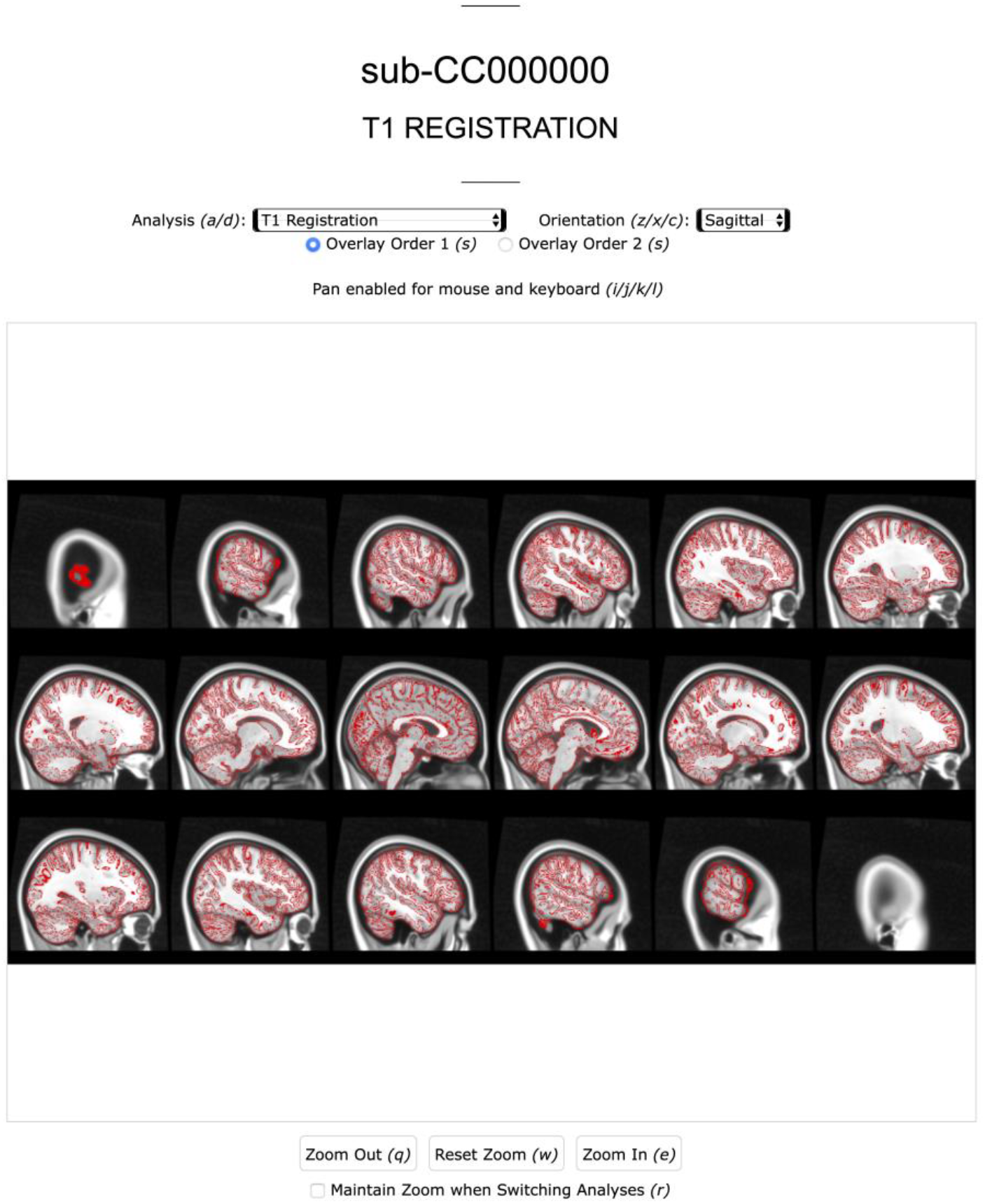
Screenshot of the Anatomical page of a subject’s QC Report. Analysis (e.g. extraction, registration (shown), segmentation) and image view can be navigated with mouse or keyboard.

As part of the QC sub-pipeline development, we included FSL’s EDDY QC toolbox for generating automated reports of within-(EDDY QUAD) and across- (EDDY SQUAD) subject QC assessments. Reports automatically generated by these tools, along with others from FEAT and MELODIC can be found in our QC Report. Notably, our QC Report reconstructs the existing MELODIC ICA report and combines it with classified ICA outputs from FIX into a single MELODIC page. This page groups signal and noise labelled components for quick assessment of FIX performance and allows immediate access to every component’s analyses through a set of dropdown menus and optional hotkeys.

The QC Report is portable, at ~180MB for a compressed QC Report compared to ~2GB to ~5GB for a compressed full subject for the datasets we have tested. This enables faster and lower-overhead report sharing and collaboration without needing to share potentially sensitive raw or intermediate data. Furthermore, viewing the report requires no installations and it can be run on any operating system and modern browser. The lightweight and portable nature of our report is especially impactful for users who work on headless servers and may need to download files for visualization.

### 2.7 The Brain Imaging Data Structure (BIDS)

During processing, we retain and mimic the directory structure and file organization of the UK Biobank pipeline. We extend the UK Biobank’s BIDS conversion script, which organizes pipeline output files in a manner outlined in a filename conversion dictionary. Our extension updates the conversion dictionary with BIDS-compliant filenames for new TVB-UKBB intermediate and output files. This ensures interoperability of our pipeline’s outputs, such that the derivative and raw data files for each subject are named, documented, and organized in a directory structure in accordance with BIDS v1.6.0. Additionally, we have introduced a reversal feature to the BIDS conversion script, allowing BIDS-converted pipeline outputs to be reverted to the original TVB-UKBB file organization to facilitate reprocessing and reproducibility.

### 2.8 Developed software

The pipeline has been constructed principally with Linux compatibility in mind. The software utilizes a Python backbone which brings together various BASH, MATLAB, and R scripts to process data moving through the pipeline. This software environment is encapsulated largely in a conda environment which can be used standalone or inside a supplied Singularity container (Kurtzer et al., 2017). The installation is straightforward and self-contained, with minimal dependencies on external applications after configuration. The Singularity container enables users to stage and run the pipeline in myriad high-performance computing environments and to leverage the batching capabilities of schedulers like SLURM and SGE.

### 2.9 GitHub repository and documentation

The source code for our pipeline is hosted on GitHub (https://github.com/McIntosh-Lab/tvb-ukbb). Several versions of the pipeline exist, each catering to different dataset needs and specifications. These versions are stored as separate branches on the repository. For example, branch Cam-CAN is available for pipeline users who want to process Cam-CAN subjects or datasets similar in specification to the Cam-CAN dataset using the Singularity container. ADNI3 is similar and is also the basis for the main branch as it is likely compatible with the widest range of datasets that the pipeline would be used with.

Extensive documentation on the TVB-UKBB pipeline is available on the Wiki page of our GitHub repository. This wiki includes information on the methodological components of the pipeline as well as installation, troubleshooting, QC interpretation, usage examples, etc.

### 2.10 Installation and Singularity container

Due to the high degree of complexity involved in the UK Biobank pipeline installation process, significant efforts were made to streamline installation and configuration. Singularity is a core component of these streamlining efforts due to its use in high performance computing environments as well as its ability to encapsulate complex and difficult-to-configure software stacks. Users may wish to install our pipeline with or without the Singularity container. All dependencies are included in the Singularity container, with the exception of FreeSurfer, AFNI, and ANTS. FSL and CUDA 9.1 were installed and configured in the container because GPU-enabled versions of BEDPOSTX, EDDY, and PROBTRACKX all require CUDA 9.1. MATLAB compatibility is packaged into the container using the MATLAB Compiled Runtime to eliminate the need for a MATLAB licence.

### 2.11 Technical features

The pipeline features CPU-only and CUDA-enabled versions. The CUDA-enabled version allows the FSL toolkit to take advantage of NVIDIA GPUs to drastically reduce runtimes of the BEDPOSTX, EDDY, and PROBTRACKX programs and cut the overall pipeline runtime significantly. If NVIDIA GPUs are not available, users can specify the CPU-only version which will run these FSL toolkits serially. To shorten the runtime and memory requirements of probabilistic tractography on CPU, we also include a parallelized implementation of PROBTRACKX.

Due to the variety of programming languages and heavy use of BASH, efforts were made to simplify configuration of pipeline parameters for end-users. The result is a single configuration file where the vast majority of environment variables for pipeline configuration and customization are specified. Parameters like the location of a FreeSurfer installation, specification of parcellation, etc. are set in this configuration file and is sourced prior to running the pipeline.

## 3 Results

### 3.1 Usage

The pipeline currently supports Cam-CAN and ADNI3 datasets, and can be customized by naive users to support novel datasets. Here we demonstrate usage of the TVB-UKBB pipeline using the Cam-CAN dataset (cite), which includes T1w, T2*, resting-state and task-fMRI, field maps, and dMRI from ~650 adults aged 18-99. In these examples, we used a Schaefer-Tian parcellation consisting of 400 cortical and 20 subcortical regions.

The key TVB inputs generated by the pipeline can be visualized and analyzed with ease. Figure 6 shows the pipeline outputs of interest for connectome-based modelling for an example subject. These include the structural connectivity weights and tract lengths matrices, and the resting-state BOLD-fMRI responses and functional connectivity matrix.

**Figure 6.**
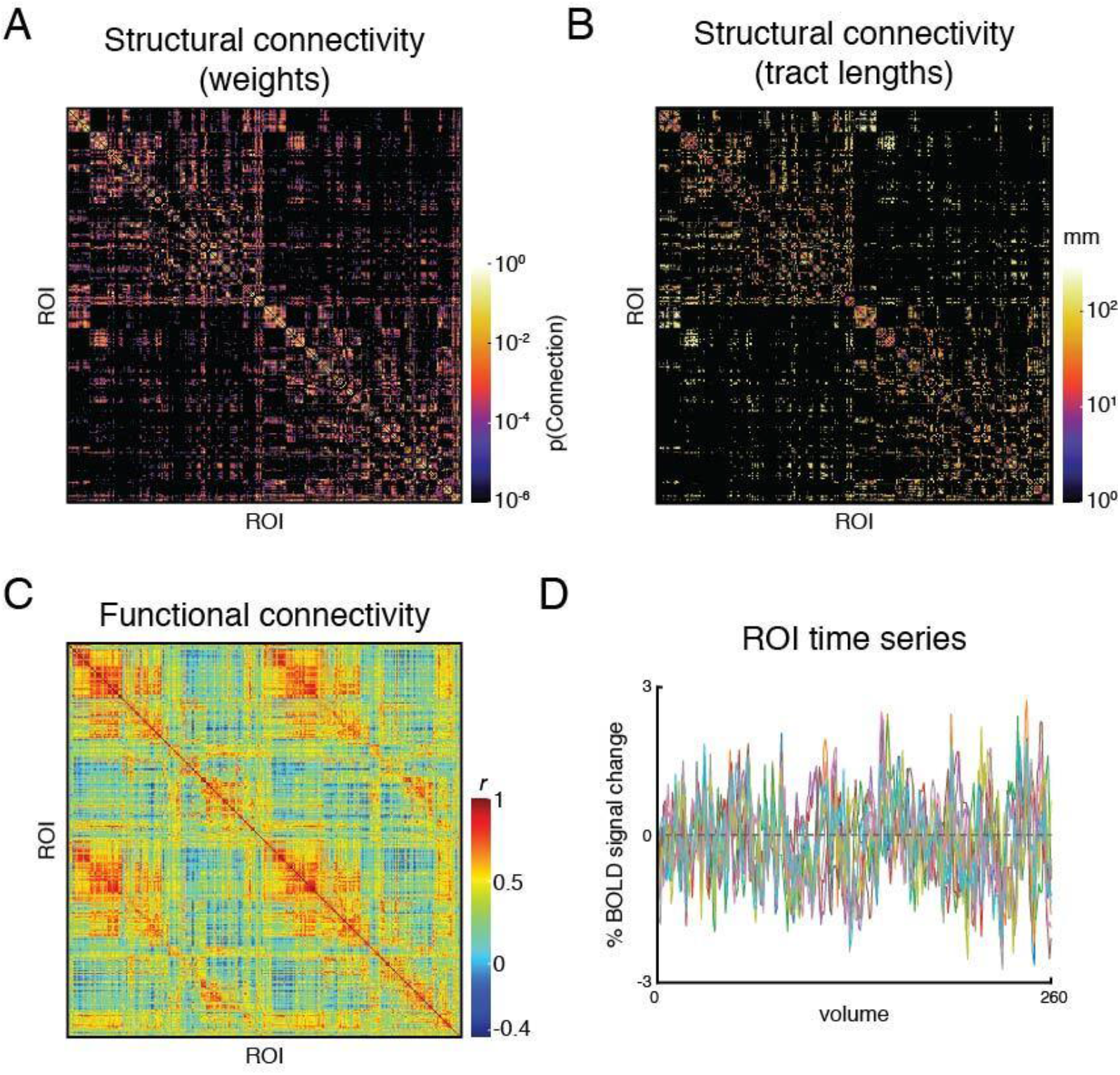
An example subject’s set of pipeline outputs for connectome-based modelling. These include (A) a weights matrix and (B) a tract lengths matrix from dMRI processing that capture the subject’s structural connectivity; (C) a functional connectivity matrix of Pearon correlation coefficients, and (D) the region of interest (ROI) time series from resting-state fMRI processing. The structural connectivity matrices are presented on a log scale to enhance readability. Ten ROIs were chosen randomly for presentation in (D).

### 3.2 QC procedures & QC Report Usage

The QC reports allow users to quickly inspect pipeline intermediates and outputs. A detailed manual QC of a single subject without the QC report previously took our experienced raters (DS, KS) up to 30 mins to complete, but a subject assessed with the QC report now takes an average of ~ 5 mins. Here we briefly outline our QC procedures for aging (Cam-CAN) and neurodegenerative (ADNI3) imaging data and provide some examples of common preprocessing errors detected using the QC reports. We describe the QC procedures in the order that the pipeline processes the data, but in practice we start QC investigations with the final outputs of the pipeline (structural and functional connectivity and functional responses) and work upstream through the QC report to quickly pinpoint the source of errors in processed subjects.

#### 3.2.1 Structural sub-pipeline QC

Examination of the structural pipeline includes the raw T1w image and the outputs of T1w brain extraction, segmentation and registration to the MNI template. The reconstructed T1w image is checked for the presence of major motion or other visible artifacts. The T1w brain mask is then inspected and inclusion of dura along the lateral boundaries is noted.

The labelled and unlabelled segmentation outputs are also examined, and the accuracy of tissue classification (especially the delineation of grey and white matter) is assessed. Misclassification of non-brain tissue (i.e., inclusion in grey and white matter segmentations) is also noted. For older adults in the Cam-CAN sample (≥50 years), we also checked if white matter lesions were misclassified as grey matter during segmentation. This was supported by also inspecting the T2* image in conjunction with the T1w. Figure 7 shows an example of white matter lesions being classified as grey matter. In cases with high WML loads, this will be impossible to avoid, and QC involves deciding to what extent the misclassification impacts tractography, namely the placement of seed and target ROIs, which will be covered below.

**Figure 7.**
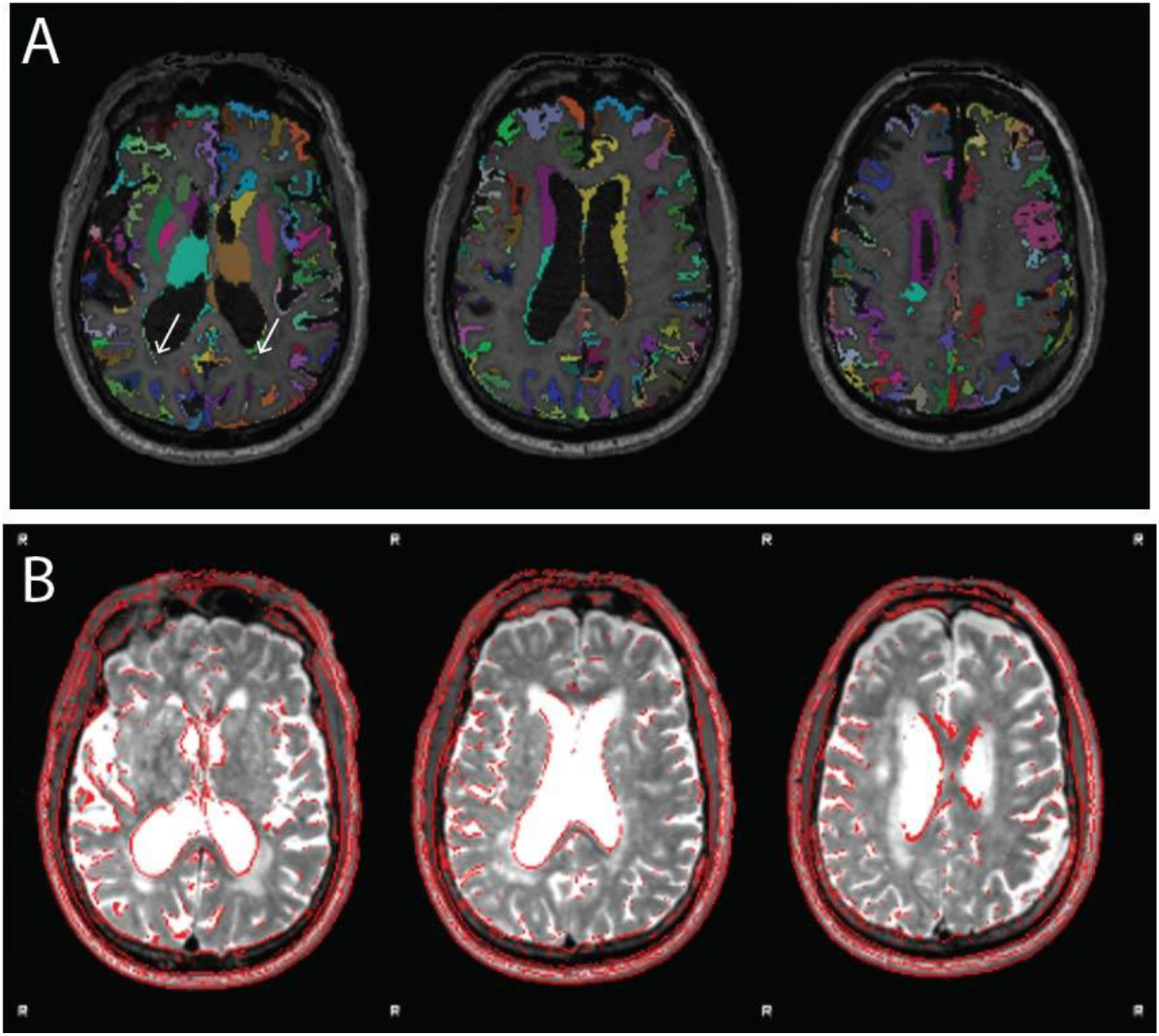
Example of white matter lesion misclassification as grey matter. (A) The labelled grey matter image is shown on the T1w. (B) T2* image from the same older adult subject indicating a significant volume of white matter lesions that are also notable on the T1w. Although performing segmentation on the T1w image using age-specific tissue priors is largely successful despite the large white matter lesion volume, some misclassification remains (white arrows in panel A). Images reproduced from the example subject’s QC report.

Finally, the registrations of the structural images to the MNI template are also inspected. Poor brain extraction and/or significant brain atrophy can affect the quality of the registration. Since the parcellation is defined on the MNI template, poor registrations can substantially hinder the parcellated downstream outputs from both the functional and diffusion sub-pipelines.

Similar procedures are followed for examining T2* images. For T2 FLAIR images, like those in the ADNI3 dataset, lesion classification outputs from BIANCA are also examined.

#### 3.2.2 Functional sub-pipeline QC

For the purposes of creating modelling inputs for TVB, we focus here on QC of the processing of resting-state fMRI data. For these data, the hyperlinked *feat* report is used to check the field map registration and correction, the relative motion of the resting-state fMRI scans and their registrations to both the T1w and MNI152 template. Signal dropout in susceptible areas such as the temporal pole or orbitofrontal cortex, if substantial, is also noted. The MELODIC page of the QC report is used to examine the components classified as signal to determine whether substantial artefactual components were included post-processing.

The functional connectivity matrix is visually inspected in the QC report and is checked for the presence of strong homotopic connectivity, clear delineation of intra- and inter-hemispheric quadrants, a sensible range of correlation values and minimal “banding” which can reflect motion artifacts or misregistration of the parcellation. The QC report allows users to examine the matrix in conjunction with a carpet plot of the cleaned ROI time series and the MCFLIRT motion plots to determine whether residual motion artifacts impact the functional connectivity matrix. See Figure 8 for an example of a bad resting-state fMRI processed outcome.

**Figure 8.**
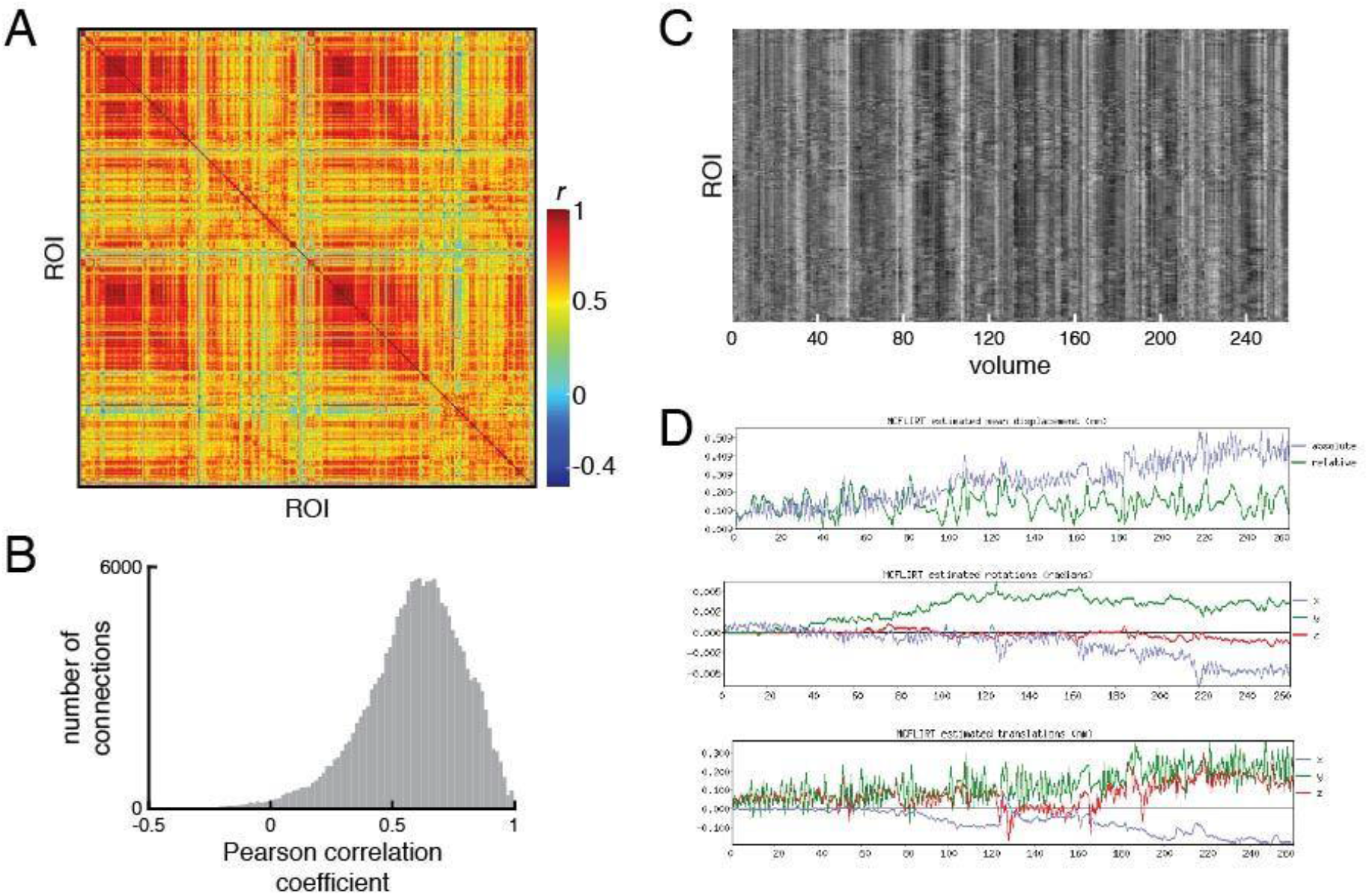
An example of poorly processed resting-state fMRI. (A) Functional connectivity matrix and (B) distribution of functional connectivity show large number of strong positive correlations and a compressed range of correlations. (C) Examination of the carpet plot of region of interest (ROI) time series suggests artefacts remain in fMRI data after cleaning. (D) In the QC Report, motion estimations from MCFLIRT are shown alongside the carpet plots for quick assessment. All images reproduced from the example subject’s QC Report.

#### 3.2.3 Diffusion sub-pipeline QC

The QC procedure for the diffusion sub-pipeline starts with examining the undistorted B0 image to check the quality of distortion correction and the presence of major artifacts. The brain mask calculated from the distortion corrected B0 is also checked as it is used to exclude non-brain tissue from downstream diffusion processing. Brain masks that are too conservative are noted as they can impact registration and placement of ROIs for tractography. The principle orientations of the modelled fibres are also inspected to confirm that the b-vectors have been specified appropriately. It is usually necessary to check the orientations for a single representative subject per study, but in the case of multi-site studies the user may wish to check representative subjects from each site. The registration between the reference B0 image and the T1w is also examined.

Next, the inputs for tractography are examined. These include the grey matter exclusion mask, and the seed and target ROIs that are overlaid on the FA image in the QC report. Each of these images are checked for accuracy of their placement. The border of the brain is also inspected and seeds that are mislocalized to dura or other non-brain tissue is noted (see Figure 9 for example of poor quality tractography seed placement). With atrophic cases, poor T1-MNI template registration can impact the quality of the tractography within the brain and those with a large white matter lesion load will have lesions labelled as grey matter which can cause similar issues.

**Figure 9.**
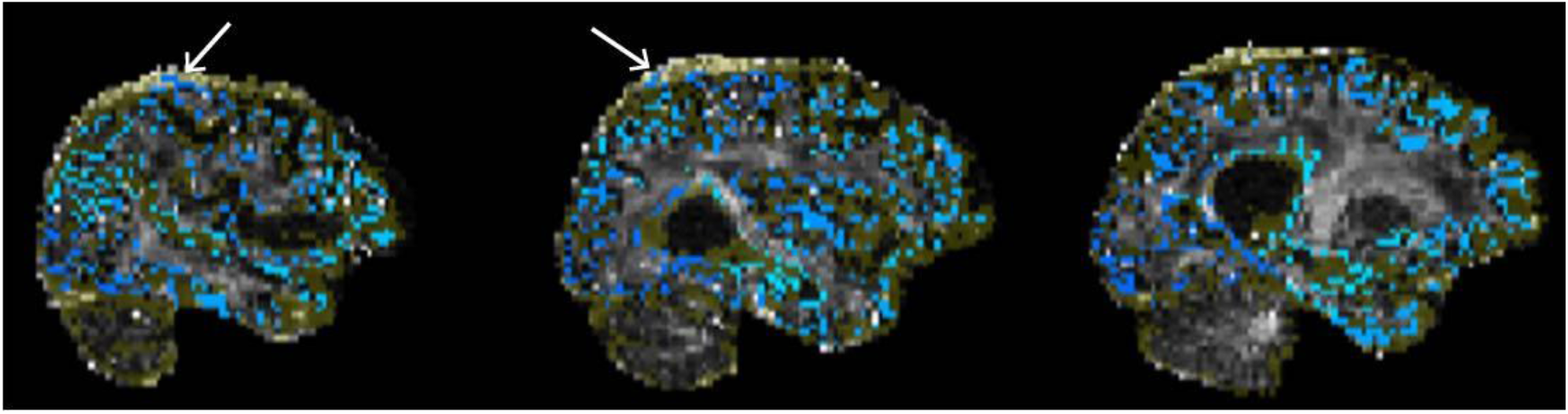
Example of poor quality tractography seed/target placement. The seeds/targets image (blue) as well as the exclusion mask image (yellow) are overlaid on the FA image. White arrows indicate seeds/targets located in the dura.

Finally, the structural connectivity matrices are examined. This includes the weights matrix, which is displayed with a logarithmic scale to improve visual assessment, and the tract lengths matrix. Visual inspection can be aided by the examination of the distributions of weights and tract lengths. Extreme sparsity of the connectome is easily detected and is often apparent in the interhemispheric quadrants of the matrices (Figure 10).

**Figure 10.**
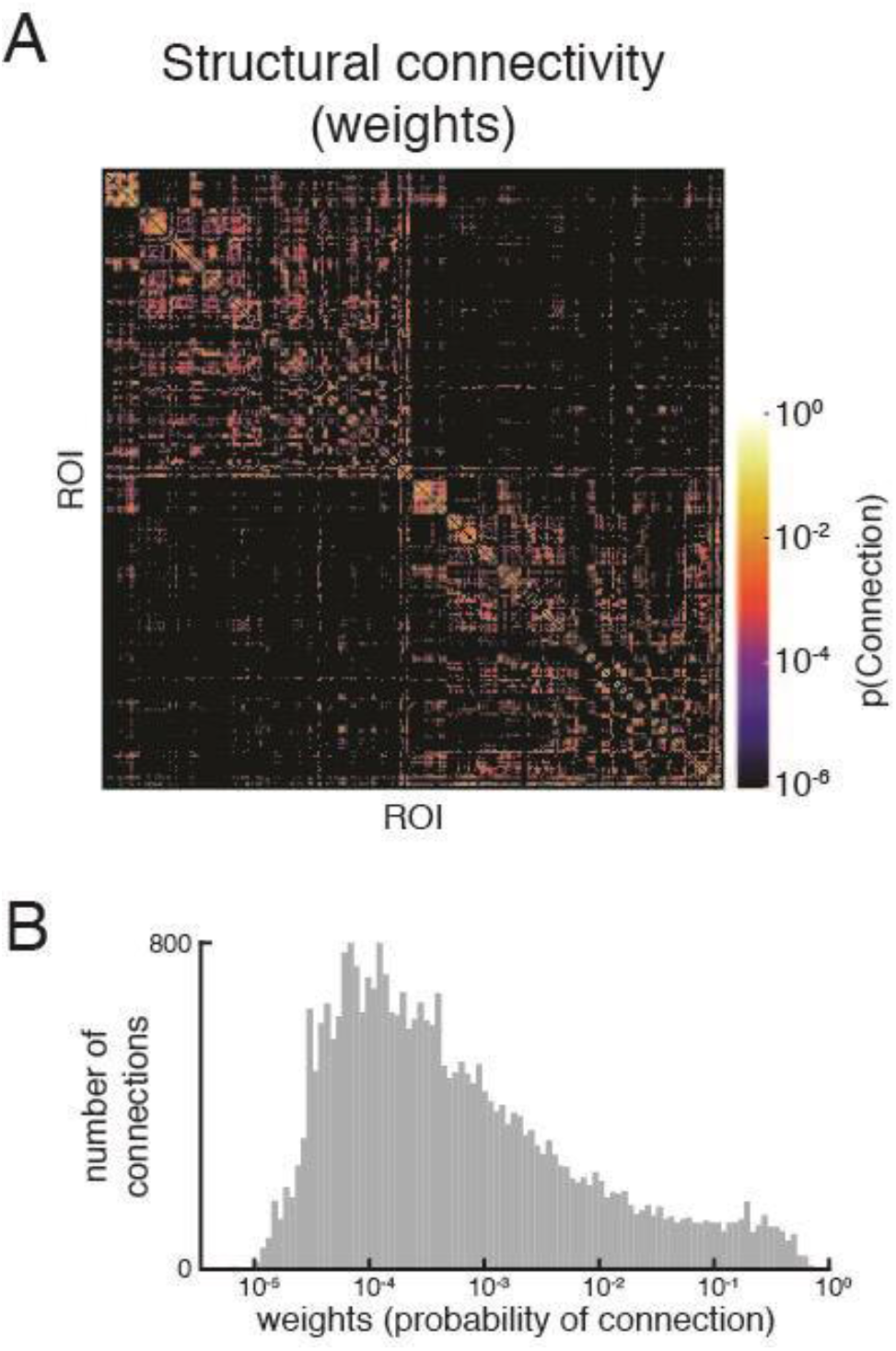
(A) An example structural connectivity matrix of poorer quality. Note the sparsity, especially in the interhemispheric quadrants (top right and bottom left), which was confirmed by (B) the relatively small distribution of non-zero weights in the matrix. Upon further examination, the dMRI registration to T1w was poor, resulting in some tractography seeds and targets being placed in non-brain tissue. Both images shown are reproduced from the QC Report.

More examples of well processed and poorly processed pipeline outputs can be found in Supplementary Figures 2 through 9.

### 3.3 Utility of new IDPs and other summary statistics

We performed manual QC of 140 Cam-CAN subjects to enable a preliminary assessment of the utility of existing and newly developed IDPs and summary statistics. This assessment was done using a partial least squares analysis of the IDPs with subjects grouped by the rater’s scores. For the functional sub-pipeline, this analysis returned one significant latent variable (Figure 11) showing how IDPs related to head motion, temporal signal-to-noise ratio, the proportion of signal/noise components, and the distribution of functional connectivity values (e.g., centre, range, shape) to be reliable indicators of resting-state fMRI processing quality (p = 0.001, 83.4% cross-block covariance).

**Figure 11.**
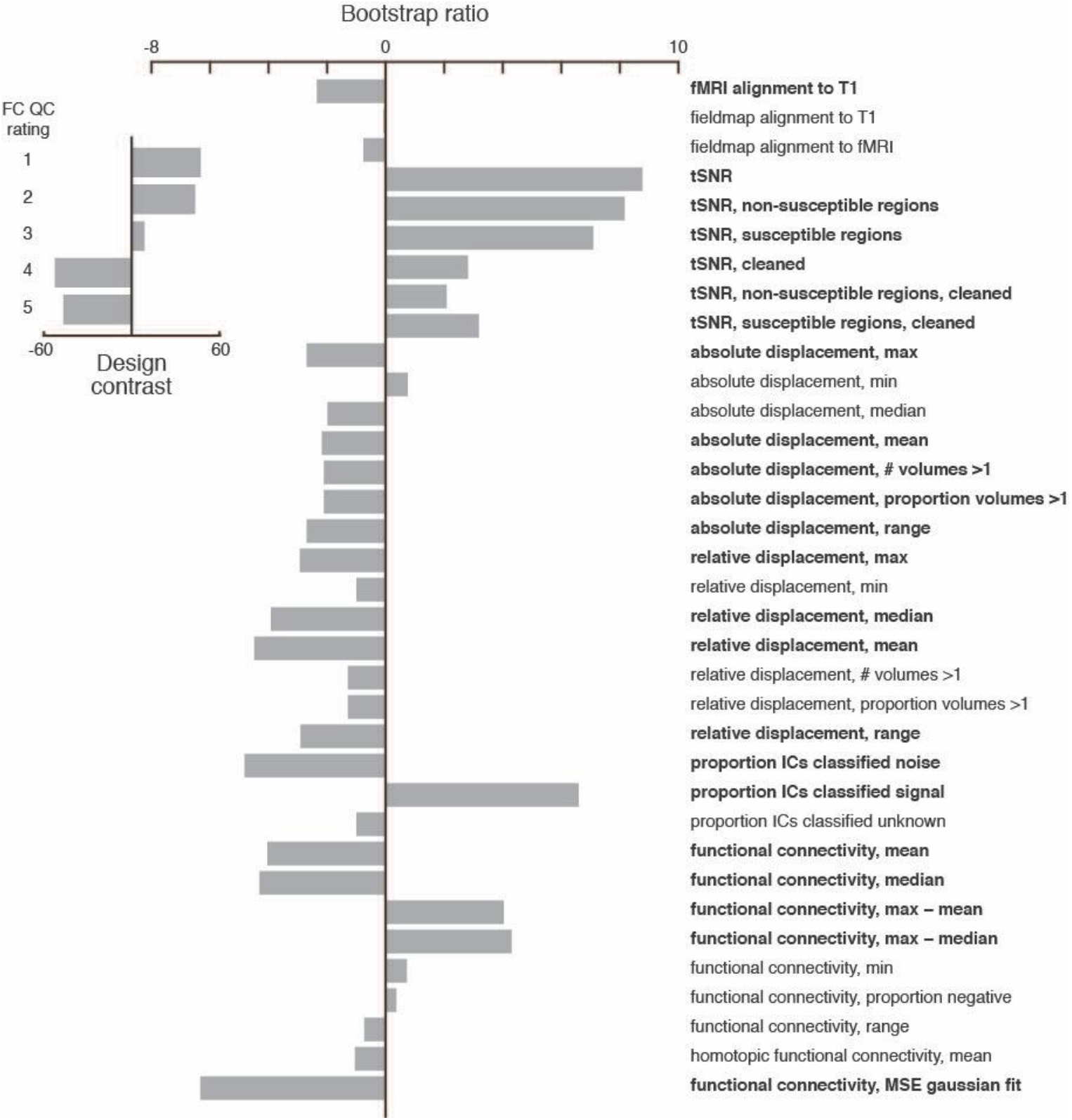
Partial least squares analysis of functional sub-pipeline IDPs and summary statistics as a function of the functional connectivity QC rating. The analysis returned a contrast (inset) between good (1 & 2) and bad (4 & 5) scores. The most reliable indicators of a good QC rating included high temporal signal-to-noise ratios, low relative displacement, a higher proportion of ICA components classified as signal, and gaussian-like functional connectivity distributions. IDPs and summary statistics with an absolute value bootstrap ratio >2 were considered reliable (see Methods), and are indicated in bold.

A similar analysis of the diffusion sub-pipeline IDPs resulted in no significant latent variables. This was likely due to a lack of variability in the quality of the diffusion processing and structural connectivity, where nearly all subjects’ (136/140) diffusion sub-pipeline outputs were judged by our raters to be either excellent (1) or very good (2).

## 4 Discussion

We have described the development of the TVB-UKBB pipeline, an open-source, easy to install, automated multimodal MRI processing solution for generating inputs for connectome-based modelling that directly interface with TheVirutalBrain. The pipeline has been containerized and supports various job schedulers on high performance compute clusters. We have tested it on both healthy and clinical populations and added features to improve its robustness against the morphological changes observed in aging and dementia.

We developed the TVB-UKBB pipeline with the processing of aging and neurodegenerative data, such as those from ADNI (Mueller et al., 2005) and Cam-CAN (Taylor et al., 2017), in mind. These datasets present particular challenges such as significant changes in brain morphology with age and/or disease (i.e., brain atrophy) and decreased image contrast, which can greatly affect registrations to a standard template and the classification of tissue classes. We addressed inaccuracies in grey matter classification by either taking advantage of available T2 FLAIR images for classifying white matter lesions, or by using age-specific tissue priors when T2 FLAIR images are not available. Future developments will include a fuller implementation of age-specific or, more generally, study-specific templates to aid registrations.

Our pipeline offers an alternative for generating modelling inputs to pipelines that rely on working with cortical surfaces. This avoids the need to project lower resolution data to high resolution surfaces (Alfaro-Almagro et al., 2018), avoids manual interventions that might be needed for correcting tissue segmentations of aging and neurodegenerative data (McCarthy et al., 2015; Henschel et al., 2020; Srinivasan et al., 2020), and avoids the long processing times needed for reconstructing the cortical surface. It also allows for easier integration of subcortical region parcels that, until very recently, were not available on the surface (but see Lewis et al., 2022). We added the ability to perform distortion correction on dMRI data for datasets without reverse phase-encoded images by adopting a toolbox that generates a synthetic undistorted B0 image (Schilling et al., 2019). Tractography methodologies for our pipeline were chosen based on our previous validation work comparing probabilistic tractographic outputs to connectomes derived from anatomical tracer data in macaques (Shen et al., 2019b). We found this method to produce reasonable estimates of fibre tract capacities (or “weights”) and fibre tract lengths. All of these considerations were made so that a greater range of ‘legacy’ datasets could be accommodated by our pipeline. Although these were all important, we recognize that cortical surface processing is considered state-of-the-art because it handles the problem of partial voluming effects and accommodates spatial smoothing to increase the signal-to-noise ratio (Brodoehl et al., 2020). Basic FreeSurfer support is already available as a part of the UK Biobank pipeline and future in-depth integrations with our pipeline are planned. GPU-enabled deep learning implementations, in particular, are attractive for creating more accurate cortical surface reconstructions quickly in aging and neurodegenerative data (Henschel et al., 2020). Given the increasing availability of GPU processing, this is in line with our efforts to develop a faster and more consistent pipeline. This type of cortical surface reconstruction will be especially important for developing M/EEG processing sub-pipelines where cortical surfaces are needed for computing the forward solution for source localization. Users may also wish to use other tractography approaches such as those that constrain tractography using anatomical priors (Smith et al., 2012). The modular implementation of our pipeline allows for these future adaptations to be implemented with relative ease.

A key component of our pipeline is the development of user-friendly HTML reports to facilitate QC assessment and faster subject scoring. With the introduction of hotkeys, fully navigable pre-generated image overlays, and re-compilation of FSL reports, our QC Reports make the novel and essential images generated by the QC sub-pipeline accessible. Existing reports are also consolidated with these images into a single, convenient point of access with an intuitive interface.

To further support QC efforts for large multimodal datasets, we developed a number of new image-based metrics and summary statistics for assessing resting-state fMRI and dMRI processing. The summary statistics, in particular, capture characteristics of processed data (i.e., connectivity matrices) that may still reflect residual artifacts that remain post-processing. For example, high motion indicated by simple motion related metrics may not warrant exclusion of a subject because some motion artifacts can be detected and removed. Post-processing summary metrics related to the FC can convey information about the successful or unsuccessful removal of motion artifacts which cannot be derived from simple motion-related metrics that are typically available in other QC reports. Image-based metrics from the UK Biobank’s structural sub-pipeline has proved useful for training a classifier to detect poorly-processed data (cite UKBB paper). Our preliminary assessment with a partial least squares analysis of our newly developed metrics suggest that extending the machine learning approach to include our new downstream metrics could be useful for automated QC.

We developed our pipeline with the FAIR principles for data (Wilkinson et al., 2016) and software (Lamprecht et al., 2019; Katz et al., 2021) management in mind. We adopt the BIDS neuroimaging standard (Gorgolewski et al., 2016) for raw data file naming, directory organization and metadata and extend the standard to the derived data. The source code is publicly available under the Apache 2.0 License, version controlled and supported by wiki-style documentation and a discussion board. Its containerization improves both accessibility and interoperability and its customization options allow for reuse across different datasets and research applications. Future iterations of the Singularity container will include FreeSurfer, AFNI, and ANTS once a solution to circumvent cloud storage quotas has been implemented.

Our pipeline generates multi-modal outputs for connectome-based modelling that are directly compatible with TheVirtualBrain software package. The high throughput nature of the pipeline, its robustness against the challenges imposed by MRI imaging of aging and clinical populations, and its extended QC capability contribute to the expanding scope of TheVirtualBrain project. In combination with the growing availability of datasets that span large age ranges and different neurological disorders, our pipeline supports TheVirtualBrain project’s endeavours to understanding large-scale network dynamics at the level of the individual.

## 6 Author Contributions

KS and ARM conceptualized the project. KS, DS, and JW developed methodology. NFL, JW, KS, and ZW contributed to software development, implementation, and testing. AS, NFL, JW, and KS performed data curation. DS, AK, AS, and KS validated research outputs. KS, AK, and JW performed statistical analysis. KS, NFL, JW, SD wrote the initial draft of this paper. All authors reviewed and edited this paper. KS and ARM supervised research activities. ARM and KS acquired financial support for the project.

## 8 Acknowledgments

This research was enabled in part by support provided by Compute Ontario (www.computeontario.ca/) and Compute Canada (www.computecanada.ca).

## 10 Data Availability Statement

Two datasets were used for pipeline development. The ADNI3 data can be found here: http://adni.loni.usc.edu/data-samples/access-data/ and the Cam-CAN data can be found here: https://www.cam-can.org/index.php?content=dataset

