## Supplementary-Material for "A robust modular automated neuroimaging pipeline for model inputs to TheVirtualBrain"

**Supplementary Table 1.** Table of novel TVB-UKBB IDPs and their descriptions. Underlined IDP categories exist in the original UK Biobank pipeline but have been modified for the TVB-UKBB pipeline. Bolded rows define the category of IDP that the following rows fall under.

| IDP name | Description |
|---|---|
| **Category: tvb_IDP_FC_distribution**[1] | **Functional connectivity summary statistics**[1] |
| FC_distribution_min_fMRI | Functional connectivity (Pearson correlation coefficient) minimum |
| FC_distribution_max_fMRI | Functional connectivity (Pearson correlation coefficient) maximum |
| FC_distribution_median_fMRI | Functional connectivity (Pearson correlation coefficient) median |
| FC_distribution_mean_fMRI | Functional connectivity (Pearson correlation coefficient) mean |
| FC_distribution_mean_to_max_fMRI | Functional connectivity (Pearson correlation coefficient) difference between mean and max |
| FC_distribution_median_to_max_fMRI | Functional connectivity (Pearson correlation coefficient) difference between median and max |
| FC_distribution_range_fMRI | Functional connectivity (Pearson correlation coefficient) range |
| FC_distribution_proportion_neg_fMRI | Functional connectivity proportion negative |
| FC_norm_MSE_fMRI | Functional connectivity - mean squared error for a Gaussian distribution fit |
| **Category: tvb_IDP_SC_distribution**[1] | **Structural connectivity summary statistics**[1] |

| | |
|---|---|
| SC_distribution_min | Structural connectivity (log of Probability of connection) minimum |
| SC_distribution_max | Structural connectivity (log of Probability of connection) maximum |
| SC_distribution_median | Structural connectivity (log of Probability of connection) median |
| SC_distribution_mean | Structural connectivity (log of Probability of connection) mean |
| SC_distribution_mean_to_max | Structural connectivity (log of Probability of connection) difference between mean and max |
| SC_distribution_median_to_max | Structural connectivity (log of Probability of connection) difference between median and max |
| SC_distribution_range | Structural connectivity (log of Probability of connection) range |
| SC_distribution_proportion_neg | Proportion of non-zero Structural connectivity (Probability of connection) values |
| SC_nan_lines | Structural connectivity – indices of ROIs with only NaN structural connectivity (probability of connection) values |
| SC_num_nan | Structural connectivity - number of ROIs all NaN connections |
| SC_lognorm_MSE[2] | Structural connectivity - mean squared error for lognorm distribution fit[2] |
| **Category: tvb_IDP_FIX_classes** | **MELODIC independent component (IC) class proportions: proportion of ICs that are signal, noise, or unknown** |
| FIX_prop_IC_unknown | Proportion of MELODIC ICs that are classified as unknown by FIX |
| FIX_prop_IC_signal | Proportion of MELODIC ICs that are classified as signal by FIX |

| | |
|---|---|
| FIX_prop_IC_noise | Proportion of MELODIC ICs that are classified as noise by FIX |
| **Category: tvb_IDP_MCFLIRT_disp** | **MCFLIRT displacement summary statistics** |
| MCFLIRT_rel_disp_min_fMRI | MCFLIRT relative displacement minimum |
| MCFLIRT_rel_disp_max_fMRI | MCFLIRT relative displacement maximum |
| MCFLIRT_rel_disp_median_fMRI | MCFLIRT relative displacement median |
| MCFLIRT_rel_disp_mean_fMRI | MCFLIRT relative displacement mean |
| MCFLIRT_rel_disp_range_fMRI | MCFLIRT relative displacement range |
| MCFLIRT_rel_disp_proportion_gt_one_fMRI | MCFLIRT relative displacement - proportion of time units with displacement greater than 1mm |
| MCFLIRT_rel_disp_num_gt_one_fMRI | MCFLIRT relative displacement - number of time units with displacement greater than 1mm |
| MCFLIRT_abs_disp_min_fMRI | MCFLIRT absolute displacement minimum |
| MCFLIRT_abs_disp_max_fMRI | MCFLIRT absolute displacement maximum |
| MCFLIRT_abs_disp_median_fMRI | MCFLIRT absolute displacement median |
| MCFLIRT_abs_disp_mean_fMRI | MCFLIRT absolute displacement mean |
| MCFLIRT_abs_disp_range_fMRI | MCFLIRT absolute displacement range |
| MCFLIRT_abs_disp_proportion_gt_one_fMRI | MCFLIRT absolute displacement - proportion of time units with displacement greater than 1mm |
| MCFLIRT_abs_disp_num_gt_one_fMRI | MCFLIRT absolute displacement - number of time units with displacement greater than 1mm |

| | |
|---|---|
| **Category:**<br>**tvb_IDP_homotopic** | **Mean Pearson correlation coefficient value for functional connectivity between homotopic ROIs.** |
| FC_homotopic_mean_fMRI | Homotopic functional connectivity mean |
| **Category:**<br>**tvb_IDP_func_TSNR** | **Temporal signal-to-noise ratios in fMRI** |
| fMRI_TSNR | Temporal signal-to-noise ratio in the pre-processed fMRI - reciprocal of median (across brain voxels) of voxelwise mean intensity divided by voxelwise timeseries standard deviation |
| fMRI_cleaned_TSNR | Temporal signal-to-noise ratio in the artefact-cleaned pre-processed fMRI - reciprocal of median (across brain voxels) of voxelwise mean intensity divided by voxelwise timeseries standard deviation |
| fMRI_num_vol | Number of volumes (timepoints) in fMRI scan |
| **Category:**<br>**tvb_IDP_func_susceptibility_SNR**[3] | **Temporal signal-to-noise ratios in susceptible and non-susceptible brain regions in fMRI**[3] |
| fMRI_non-susceptible_TSNR | Temporal signal-to-noise ratio in the minimally-processed fMRI non-susceptible regions - median (across brain voxels) of voxelwise mean intensity divided by voxelwise timeseries standard deviation |
| fMRI_non-susceptible_cleaned_TSNR | Temporal signal-to-noise ratio in the artefact-cleaned pre-processed fMRI non-susceptible regions - median (across brain voxels) of voxelwise mean intensity divided by voxelwise timeseries standard deviation |
| fMRI_susceptible_TSNR | Temporal signal-to-noise ratio in the minimally-processed fMRI susceptible regions - median (across brain voxels) of voxelwise mean intensity divided by voxelwise timeseries standard deviation |
| fMRI_susceptible_cleaned_TSNR | Temporal signal-to-noise ratio in the artefact-cleaned pre-processed fMRI susceptible regions - median (across brain voxels) of voxelwise mean intensity divided by voxelwise timeseries standard deviation |

| Category: tvb_IDP_all_align_to_T1 | Discrepancy between various modalities registered to T1 space and the T1 image |
|---|---|
| T2_FLAIR_align_to_T1 | Discrepancy between the T2_FLAIR brain image (linearly-aligned to the T1) and the T1 brain image |
| dMRI_align_to_T1 | Discrepancy between the dMRI brain image (linearly-aligned to the T1) and the T1 brain image |
| fMRI_align_to_T1 | Discrepancy between the fMRI brain image (linearly-aligned to the T1) and the T1 brain image |
| fMRI_fieldmap_align_to_T1 | Discrepancy between the fMRI gradient echo field map brain image (linearly-aligned to the T1) and the T1 brain image |
| **Category: tvb_IDP_fieldmap_func_align** | **Discrepancy between the fMRI field map brain image registered to fMRI func space and the fMRI func image** |
| fMRI_fieldmap_func_align | Discrepancy between the fMRI gradient echo field map brain image and the fMRI image |

[1] *Summary statistics for connectivity matrices (structural, functional) are calculated without the values from the main diagonal in order to exclude reflexive connectivity.*

[2] *Structural connectivity weights were fit to a lognormal distribution based on our previous observations that the heavily skewed distributions of anatomical tract tracing weights are well captured by the tractographic methodology used in the TVB-UKBB pipeline (Shen et al., 2019 Sci Data)*

[3] *Areas susceptible to fMRI dropout are defined by the user with an ROI mask image that is compatible with the user-defined parcellation (i.e., some subset of the parcellation). In our Cam-CAN usage example, this mask includes ROIs near large sinuses (orbitofrontal cortex, inferior temporal cortex, temporal pole).*

**Supplementary Table 2.** Table of anatomical QC Report features. QC Report Features are listed in order of appearance in the QC Report. Each QC Report Feature is presented in the QC Report in the order their described images and outputs are processed by the pipeline.

| QC Report Feature | Description |
|---|---|
| T1 registration to standard | Overlay of T1 and MNI T1 standard |
| T2 registration to T1 | Overlay of T1 and T2 |
| T1 brain extraction | Overlay of T1 and T1 brain mask |
| T1 WM segmentation | Overlay of T1 and WM |
| T1 GM segmentation | Overlay of T1 and GM |
| Unlabelled subcortex | Zoomed-in overlay of T1 and T1 brain mask |
| Labelled subcortex | Zoomed-in overlay of T1 and GM ROI parcellations |
| Labelled cortex | Overlay of T1 and GM ROI parcellations. |
| T2 registration | Overlay of T2 and T1. |
| T2 FLAIR BIANCA | Overlay of T2 FLAIR lesions and T2. |

**Supplementary Table 3.** Table of Diffusion QC Report features. QC Report Features are listed in order of appearance in the QC Report. Each QC Report Feature is presented in the QC Report in the order their described images and outputs are processed by the pipeline. EDDY QUAD and EDDY SQUAD reports are also included in the TVB-UKBB QC Report.

| QC Report Feature | Description |
| --- | --- |
| DTI Extraction | Overlay of B0 and brain mask. |
| DTI Tensor Orientation | Overlay of DTI fractional anisotropy (FA) and DTI V1 principal diffusion tensors. |
| DTI Registration | Overlay of T1 and DTI warped to T1 space. |
| SynB0 Warping | Overlay of DWI_B0 and b0_u |
| Tractography | Overlay of DTI FA and exclude, WMGM interface, or labelled WMGM interface |

**Supplemental Table 4.** Table of SC/FC QC Report features. QC Report Features are listed in order of appearance in the QC Report. Each QC Report Feature is presented in the QC Report in the order their described images and outputs are processed by the pipeline.

| QC Report Feature | Description |
|---|---|
| Structural Connectivity | Heatmap matrix of structural connectivity (probability of connection or ‘weights'). Accompanied by histogram of structural connectivity values. |
| Functional Connectivity | Heatmap matrix of functional connectivity (Pearson correlation coefficients). Accompanied by histogram of functional connectivity values. |
| ROI Carpet Plot | Carpet plot of resting-state fMRI time series for all ROIs |
| Tract Lengths | Heatmap matrix of tract lengths (‘distance’). Accompanied by histogram of tract length values. |
| rfMRI plots | MCFLIRT rfMRI estimated motion graph |
| tfMRI plots | MCFLIRT tfMRI estimated motion graph |

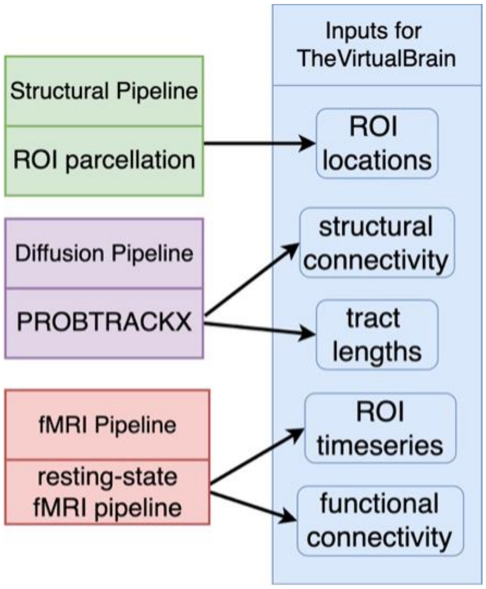


**Supplementary Figure 1**. Flow of outputs from the TVB-UKBB pipeline to the inputs used with TheVirtualBrain. Model inputs are generated by the pipeline and are compressed into a .zip file that is easily readable by TheVirtualBrain.

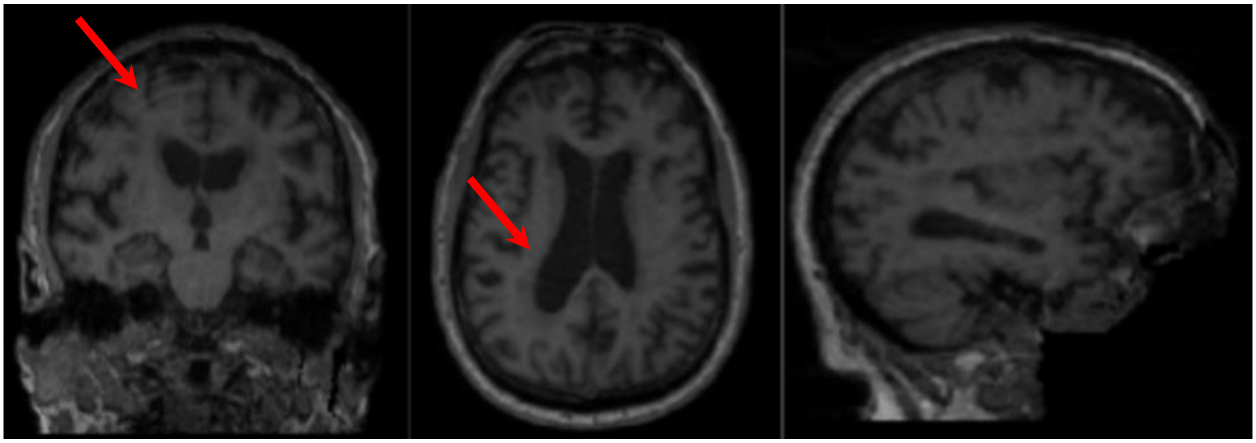

**Supplementary Figure 2.** T1 image analysis from the QC Report. Image shows a coronal, axial, and sagittal view of a subject T1 image (grayscale). Motion artifacts and atrophy are evident (red arrows).

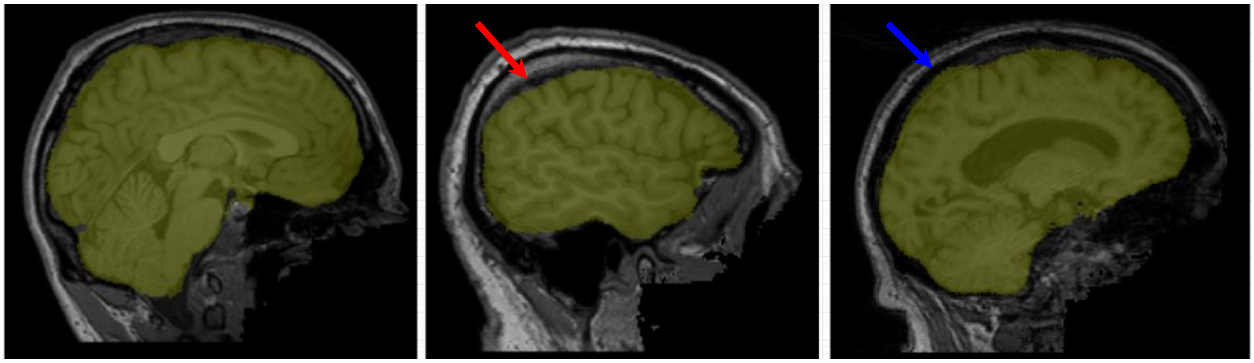

**Supplementary Figure 3.** T1 brain extraction analysis from the QC Report. Images show a sagittal view of a subject's brain mask (yellow) overlaid on top of their T1w image (grayscale). A slice from a well-processed subject can be seen on the left. A slice from a subject with insufficient brain inclusion in the mask can be seen in the middle (red arrow). A slice from a subject with excess dura mater inclusion in the mask can be seen on the right (light blue arrow).

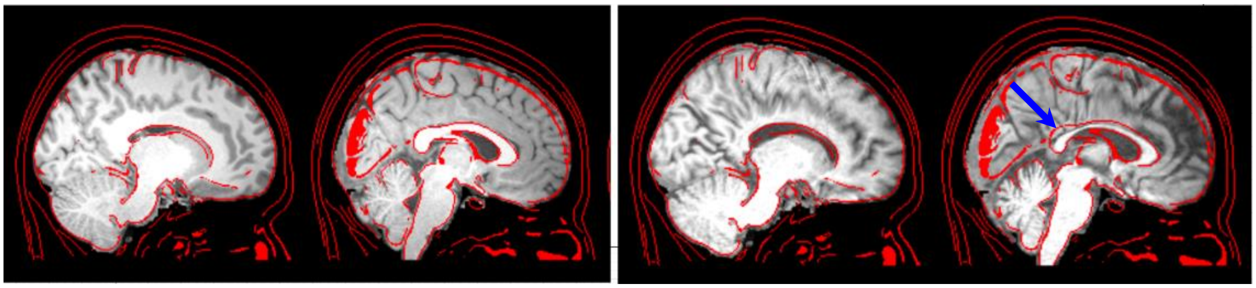

**Supplementary Figure 4.** T1 registration analysis from the QC Report. Images show a sagittal view of the MNI standard (red) overlaid on top of a subject's T1w image (grayscale). Two slices from a well-processed subject can be seen on the left and two slices from a poorly-processed subject can be seen on the right. The arrow (blue) points to an instance of poor alignment.

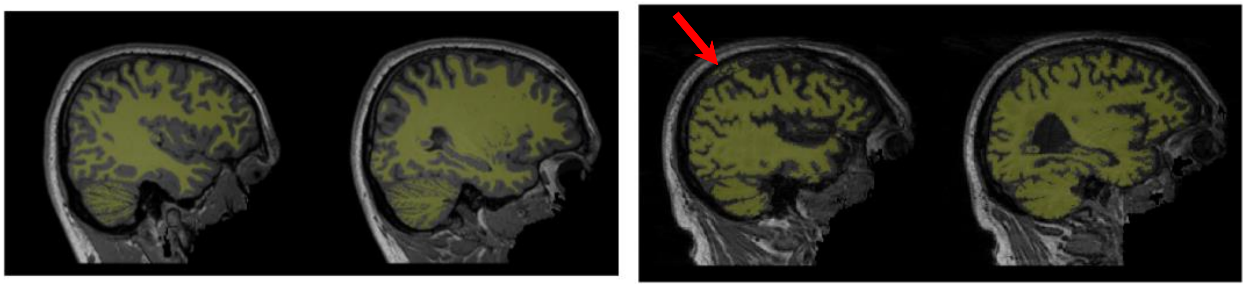

**Supplementary Figure 5.** T1 White Matter Segmentation from the QC Report. Images show a sagittal view of a subject's white matter mask (yellow) overlaid on top of their T1w image (grayscale). Two slices from a well-processed subject can be seen on the left. Two slices from a subject with misclassified white matter from the dura can be seen on the right (red arrow).

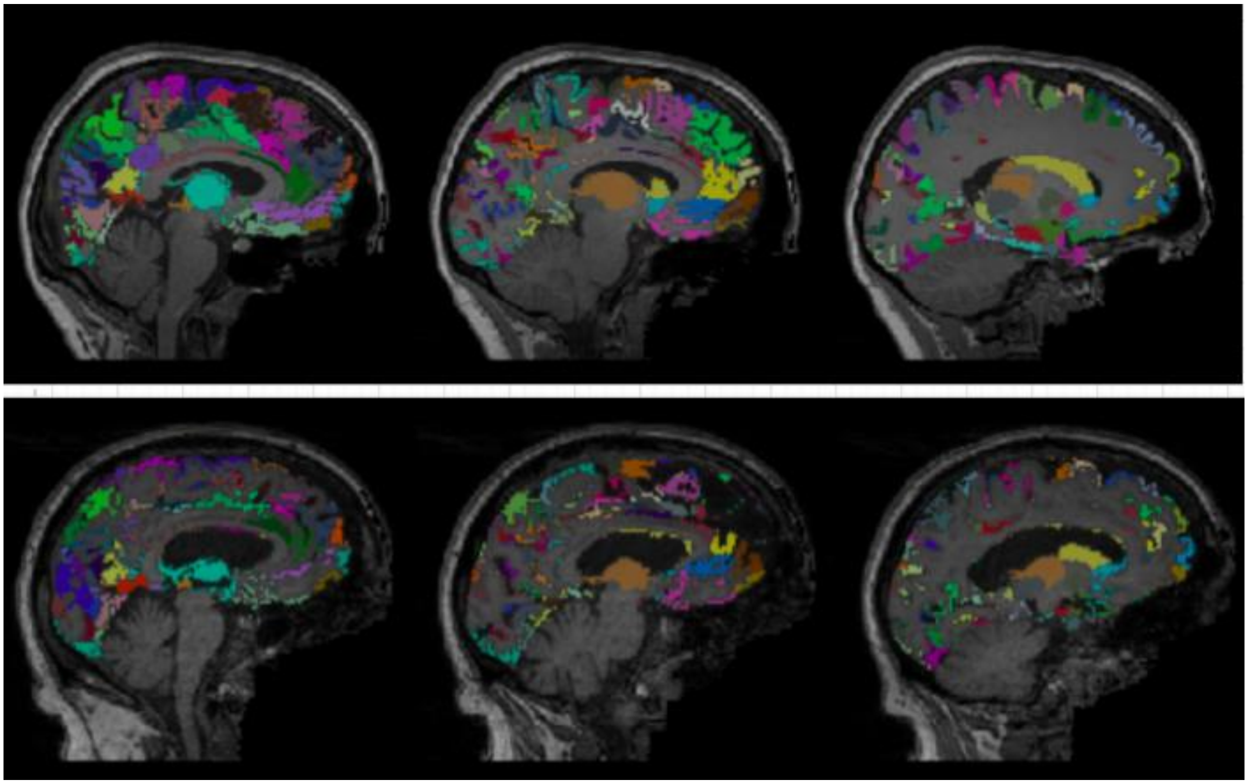

**Supplementary Figure 6.** T1 ROI segmentation analysis from the QC Report. Images show a view of a subject's labelled grey matter ROIs (coloured) overlaid on top of their T1w image (grayscale). Three slices from a well-processed subject can be seen on top and three slices from a poorly-processed subject, where grey matter labelling is sparse, can be seen on bottom.

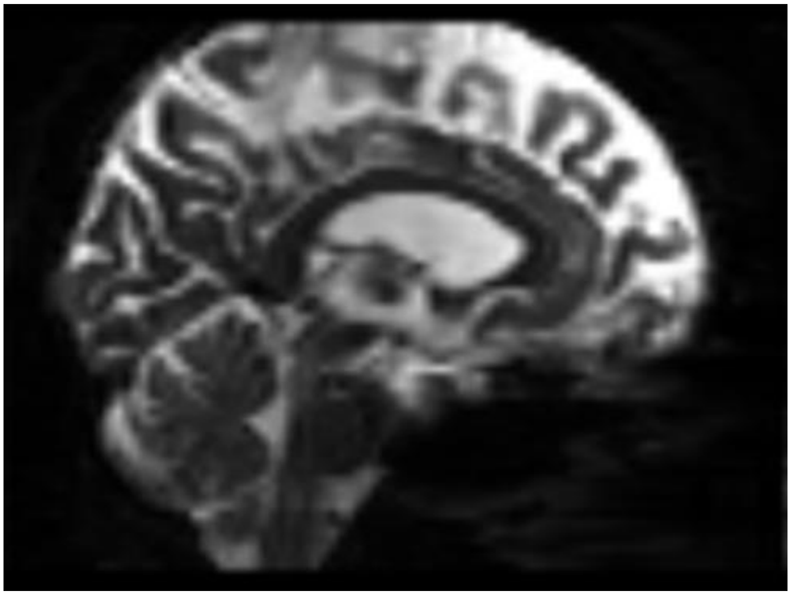

**Supplementary Figure 7.** dMRI B0 image analysis from the QC Report. Image shows a sagittal view of a subject's dMRI B0 image. The superior margins of the brain were outside the field of view.

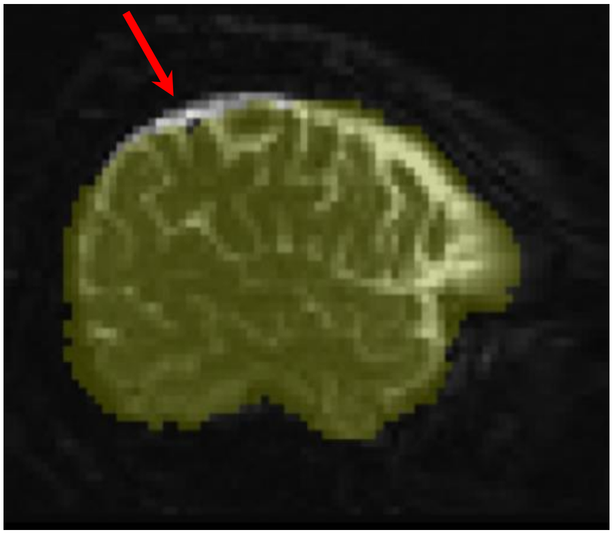

**Supplementary Figure 8.** dMRI brain extraction analysis from the QC Report. Image shows a sagittal view of a subject's dMRI brain mask (yellow) overlaid on top of their B0 image (grayscale). Insufficient brain inclusion in the mask excludes cortex (red arrow).

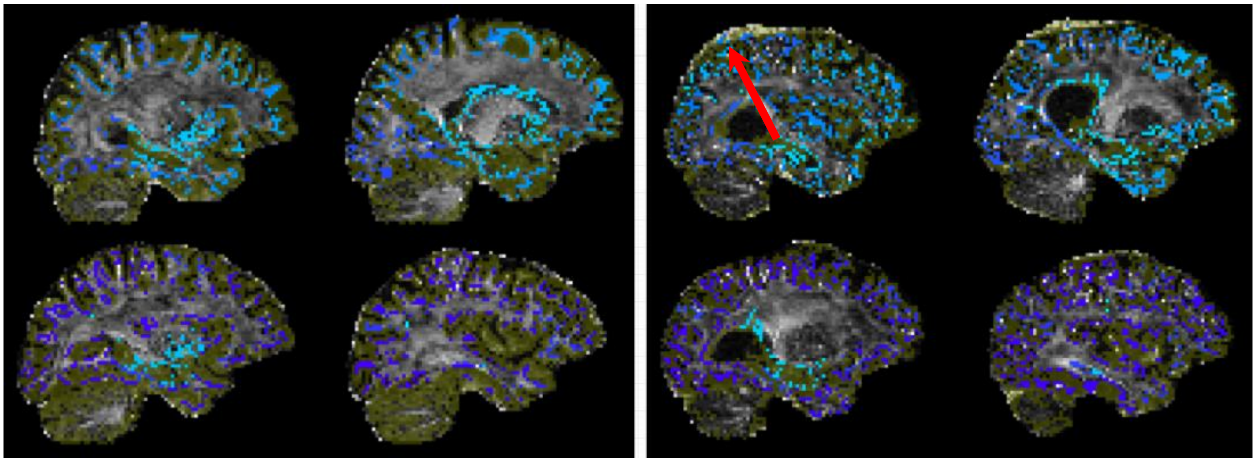

**Supplementary Figure 9.** dMRI tractography analysis from the QC Report. Image shows a sagittal view of a subject's tractography seeds (blue) and exclude mask (yellow) overlaid on top of their FA image. Slices from a well-processed subject (left) have seeds localized within the border of the brain. Slices from a poorly-processed subject (right) show seeds in the dura (red arrow).
